# Presynaptic FMRP and local protein synthesis support structural and functional plasticity of glutamatergic axon terminals

**DOI:** 10.1101/2021.06.07.447454

**Authors:** Hannah R. Monday, Shivani C. Kharod, Young J. Yoon, Robert H. Singer, Pablo E. Castillo

## Abstract

Learning and memory critically rely on long-lasting, synapse-specific modifications. While postsynaptic forms of plasticity typically require local protein synthesis, whether and how local protein synthesis contributes to presynaptic changes remains unclear. Here, we examined the hippocampal mossy fiber (MF)-CA3 synapse which expresses both structural and functional presynaptic plasticity. We report that MF boutons synthesize protein locally and contain ribosomes. Long-term potentiation of MF-CA3 synaptic transmission (MF-LTP) was associated with translation-dependent enlargement of MF boutons. Moreover, increasing *in vitro* and *in vivo* MF activity enhanced protein synthesis in MFs. Remarkably, deletion of presynaptic Fragile X mental retardation protein (FMRP), an RNA-binding protein expressed in MF boutons and previously implicated in local postsynaptic protein synthesis-dependent plasticity, blocked structural and functional MF-LTP, suggesting that FMRP is a critical regulator of presynaptic function. Thus, presynaptic FMRP and protein synthesis dynamically control presynaptic structure and function in the mature brain.

**Highlights:**

- Mossy fiber boutons (MFBs) synthesize protein locally and contain ribosomes
- Local presynaptic translation is increased by in vitro and in vivo GC activity
- MFB structural plasticity relies on de novo protein synthesis.
- Presynaptic FMRP is required for MF-CA3 structural and functional plasticity

**In Brief:** Monday et al. report that FMRP and protein synthesis in hippocampal mossy fiber boutons mediate functional and structural presynaptic plasticity.

## INTRODUCTION

Protein synthesis plays a critical role in long-term memory formation (Costa-Mattioli et al., 2009; Mayford et al., 2012) by supporting enduring structural changes at synapses (Bailey et al., 2015; Sutton and Schuman, 2006). Synaptic plasticity can result from activity-dependent changes in either or both pre- and postsynaptic compartments. Many forms of postsynaptically-expressed plasticity require local protein synthesis (Holt et al., 2019). Previous research suggests that certain forms of long-term presynaptic plasticity may rely on local protein synthesis (Hafner et al., 2019; Yin et al., 2006; Younts et al., 2016), but the molecular functions of locally synthesized proteins in axons and presynaptic compartments remain unclear. Emerging evidence suggests that dysregulation of synaptic protein synthesis may be a convergent feature of Autism Spectrum Disorders (ASDs) (Bourgeron, 2015; Klein et al., 2016; Louros and Osterweil, 2016), including Fragile X Syndrome (FXS) (Bagni and Zukin, 2019; Darnell and Klann, 2013). However, regulatory mechanisms of local presynaptic translation are poorly understood.

To explore the role of local translation in presynaptic plasticity, we took advantage of the mossy fiber (MF) axons in the hippocampus as a model system. The MF axons extend from granule cells (GC) in the dentate gyrus (DG) and project through the *stratum lucidum* to the CA3 region, where they synapse on CA3 pyramidal cells and interneurons (Acsady et al., 1998; Evstratova and Toth, 2014; Henze et al., 2000; Nicoll and Schmitz, 2005). MFs form a distinct bundle of unmyelinated fibers that form giant synapses onto the thorny excrescences protruding from proximal dendrites of CA3 pyramidal cells (MF-CA3) (Acsady *et al*., 1998; Claiborne et al., 1990) and smaller, filopodial and *en passant* synapses onto local interneurons (Acsady *et al*., 1998; Lawrence et al., 2004). MF-CA3 synapses undergo presynaptically-expressed, NMDA receptor-independent long-term potentiation (LTP) and long-term depression (LTD) (Nicoll and Schmitz, 2005). As the primary excitatory drive to the hippocampus proper, MFs are critical for hippocampus-dependent forms of learning and memory (Acsady and Kali, 2007; Rolls, 2013). Mechanistically similar forms of LTP/LTD have been identified in other brain areas (Castillo, 2012; Monday et al., 2018). Notably, MFs undergo dynamic remodeling under conditions of strong activity (Chierzi et al., 2012; De Paola et al., 2003; Maruo et al., 2016; Zhao et al., 2012), including epilepsy (Danzer et al., 2010; Sutula and Dudek, 2007), enriched environment (Galimberti et al., 2006; Gogolla et al., 2009), and spatial learning (Routtenberg, 2010).

The mechanisms by which activity leads to a presynaptic structural change are poorly understood, although a requirement for protein synthesis has been suggested (De Paola *et al*., 2003). Importantly, translation is required for MF-LTP, but whether protein is made locally or trafficked from the soma is unclear (Barnes et al., 2010; Calixto et al., 2003). Though local presynaptic protein synthesis has been suggested at this synapse (Huang and Hsu, 2004; Huang et al., 1994), the idea remains controversial (Barnes *et al*., 2010; Calixto *et al*., 2003) and it has never been directly demonstrated.

Fragile X Mental retardation protein (FMRP) is an RNA-binding protein with well-characterized roles in mRNA localization and translation (Bagni and Zukin, 2019; Darnell and Klann, 2013). Intriguingly, FMRP has been identified in a subset of presynaptic terminals throughout the brain (Christie et al., 2009), including the MF synapse. A loss-of-function mutation in *Fmr1* causes FXS, the most common monogenic cause of syndromic ASD (Hagerman et al., 2011). Under basal conditions, FMRP interacts with polyribosomes and mRNAs to form a granule (Brown et al., 2001), which can be trafficked along dendrites in an activity-dependent manner (Antar and Bassell, 2003; Darnell et al., 2011; Dictenberg et al., 2008). Dephosphorylation of FMRP by protein phosphatase 2A (PP2A) triggers granule disassembly and de-repression of translation (Narayanan et al., 2007). Regulation of dendritic mRNA by FMRP is known to regulate postsynaptic forms of plasticity (Bagni and Zukin, 2019). However, its function at the presynapse has not been well-defined, although it has been suggested to regulate translation (Akins et al., 2017; Chyung et al., 2018).

Here, we demonstrate the capacity of mossy fiber boutons (MFBs) to synthesize protein locally using cell-specific expression of a β-actin translational reporter in acute hippocampal slices. We discovered that MFBs contain ribosomes and that MF-LTP led to increases in local presynaptic actin synthesis in MFBs. Because newly synthesized actin could be related to structural plasticity at MFBs, we examined structural changes followed by MF-LTP induction. We demonstrate rapid, translation-dependent remodeling of MFBs associated with MF-LTP. Next, using a conditional KO strategy, we showed that loss of presynaptic FMRP impaired structural and functional MF-LTP, and increased local protein synthesis in the MF tract. Exposing animals to enriched environment (EE), a manipulation expected to increase GC activity, led to increased overall protein synthesis in the MF tract and specific increase of actin synthesis in MF filopodia, but did not rescue changes in protein synthesis induced by loss of presynaptic FMRP. Together, our findings provide strong evidence for the dynamic regulation of presynaptic structure and function by presynaptic FMRP and protein synthesis in the mature brain.

## RESULTS

### MFs synthesize protein locally

To test whether protein synthesis can occur in MFBs, we focused on β-actin. β-actin mRNA is highly abundant in the neuropil (Buxbaum et al., 2014; Cajigas et al., 2012) where its activity-induced translation could underlie structural plasticity of dendritic spines (Yoon et al., 2016) and growth cone guidance and remodeling (Leung et al., 2006). Given that mature MFBs are structurally dynamic (Galimberti *et al*., 2006), we examined whether actin can be synthesized locally in MFBs by optimizing a previously described Halo-actin protein synthesis reporter system for use in the mouse brain (Yoon *et al*., 2016). This system expresses β-actin protein fused to the Halotag, along with the endogenous 3’ UTR of the β-actin mRNA to allow normal mRNA localization. The bath application of ultra-bright, membrane-permeable JF dyes conjugated to the Halotag ligand (JF-HTL) allow for labeling of the Halotag fusion proteins with very high affinity and specificity (Grimm et al., 2015). We stereotactically injected a high titer lentivirus of the Halo-actin construct into the DG, which resulted in specific and sparse labeling of GCs (**Figure 1A**). Newly synthesized actin was detected using a pulse-chase experiment as follows: Bath application of the first dye (i.e. JF549-HTL) binds the entire population of pre-existing Halo-actin with an essentially irreversible covalent bond. Next, the excess dye is washed out for 30 minutes followed by the second dye (i.e. JF646-HTL) which is bath applied to label only the Halo-actin that has been newly synthesized in the intervening time window (~1.5 hours) (**Figure 1B**). To test that our approach was optimal for specifically measuring newly synthesized protein, we applied the protein synthesis inhibitor cycloheximide (CHX) during the 1^st^ wash and second dye application. CHX significantly attenuated JF646 levels in MFBs indicating that JF646 is selectively binding and reporting the levels of newly synthesized actin. To test whether Halo-actin protein could be synthesized locally, we combined two complementary approaches. First, we transected slices, making a small cut to sever the GC somas from the MF axon to prevent any trafficking of proteins from the GC soma to the MFBs (see **Supp. Figure 1A** for representative image), followed by sequential labeling of the Halo-actin. We found that MFBs were still able to synthesize Halo-actin at the same level as nontransected slices (**Figure 1C**), and there was no difference in the ratio of pre-existing to newly synthesized actin in transected slices (**Supp. Figure 1B**) suggesting that under our experimental conditions local protein synthesis is the primary contributor to the newly synthesized actin in MFBs. Importantly, reversing the order of the dye had no effect on the detection of newly synthesized proteins (**Supp. Figure 1C**). Together, these data strongly suggest that Halo-actin protein can be synthesized locally in MFBs.

**Figure 1.**
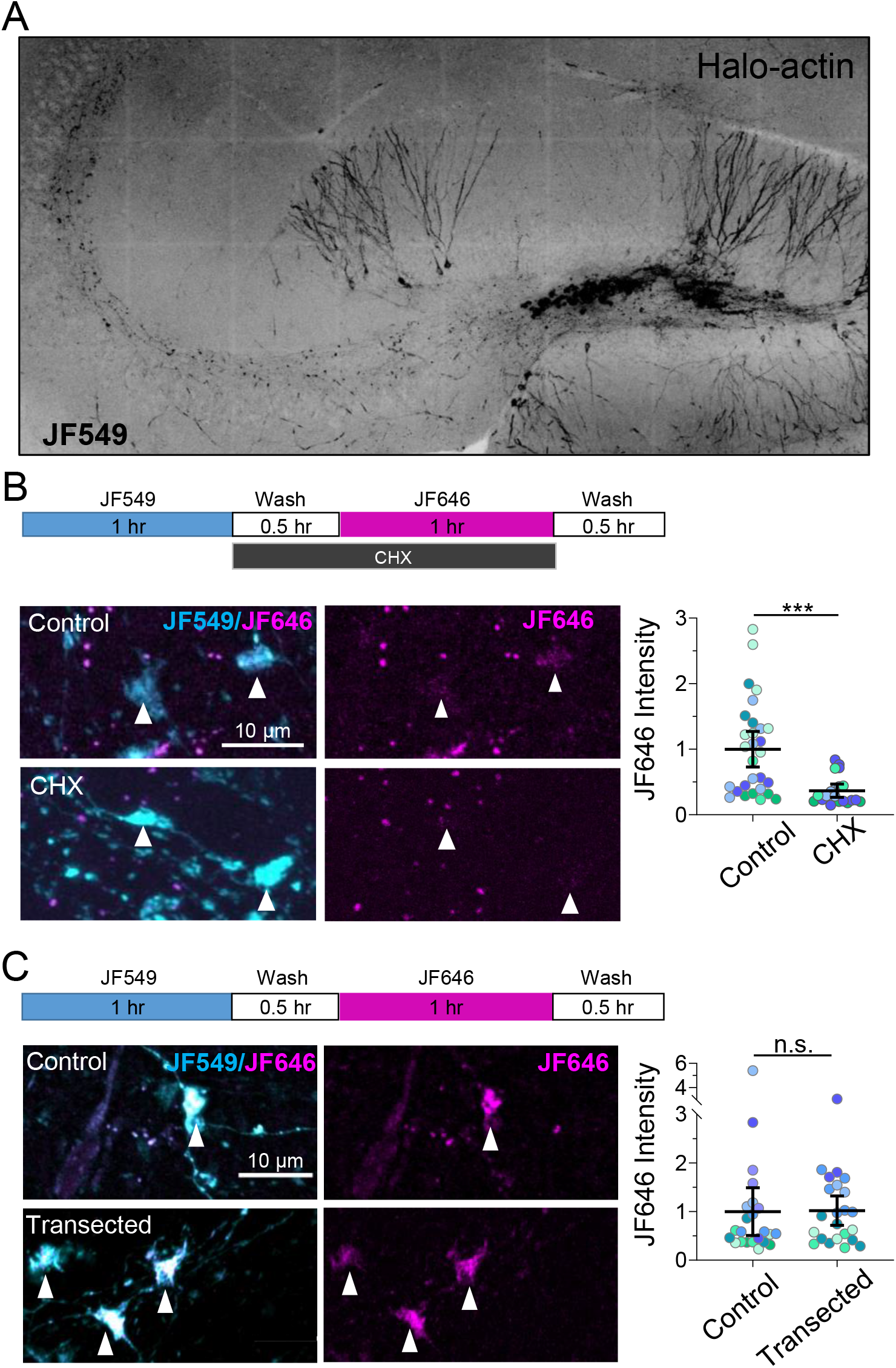
MFs synthesize protein locally. A. Representative widefield image of hippocampal slice with Halo-actin lentivirus expressing in the dentate gyrus. Halo-actin is labeled with JF549 dye. B. (*Top*) Timeline of Halo-actin pulse-chase experiment. (*bottom left*) Representative high-res Airyscan confocal images of pulse-chase labeled mossy fiber boutons in Halo-actin injected slices from transected and control slices. (*bottom right*) Mossy fiber axons that have been severed from their soma synthesized β-actin protein at the same levels as intact axons. Control: 1.0 ± 0.24 v. transected: 1.02 ± 0.15 (Mean ± S.E.M.), Mann-Whitney, U = 230, p = 0.46; n = 23 boutons, 8,7 slices respectively, 4 animals. For all Superplots in this figure, black line and bar represent the mean ± 95 % confidence interval. Points representing individual boutons are color-coded by slice and normalized to mean of Control. A. CHX bath application during second pulse significantly reduces halo-actin intensity in single MF boutons. Control: 1.0 ± 0.13 v. CHX: 0.367 ± 0.05 (Mean ± S.E.M.), Mann-Whitney, U = 94, p < 0.0001; n = 29, 20 boutons; 8, 4 slices respectively; 4 animals.

To assess local protein synthesis with high temporal resolution in nontransected MF axons, we took advantage of a newly developed photoactivatable (PA) Halo dye (Grimm et al., 2016). We bath-applied JF549-HTL and PA-JF646-HTL using the same pulse-chase paradigm as described above except after washing out excess PA-JF646, and then we photoactivated (1 s, 25 mW, 40x obj., 0.8 NA) specifically in the CA3b region where our recordings and structural analysis are performed (see below), to capture only newly made Halo-actin in MFBs (**Figure 2A**). We found that photoactivation revealed significantly more Halo-actin in MFBs than in control slices that still received the PA-JF646 dye, but no photoactivation (**Figure 2B**). These data are consistent with newly synthesized Halo-actin at MFBs. However, we cannot exclude fast axonal trafficking of new Halo-actin from the GC soma, a possibility that seems unlikely in a one-hour time window given the estimated axonal transport rates of actin (2-8 mm/day) (Roy, 2020) –e.g. a protein synthesized in the GC soma would take 3-12 hours to reach MFBs in CA3b approximately 1 mm away. To test whether Halo-actin can be trafficked to MFBs in that time frame, we performed additional experiments wherein we photoactivated Halo-actin in GCs and measured the fluorescence intensity of PA-JF646 at two time windows (1 hour and 3.5 hours) following photoactivation, where the wash time was the same between both groups. We identified boutons in CA3 using an antibody against the Halotag. Measuring PA-JF646 fluorescence in MFBs in the CA3b region showed that PA-JF646 labeled Halo-actin at 1 hour was not present above the levels in control slices with no photoactivation (**Figure 2C**), whereas we began to detect Halo-actin in this area after 3.5 hours as expected (**Figure 2D**). Summary data showed significant difference in all three CA3 subregions between the two timepoints (**Figure 2E,F**). These data indicate that Halo-actin can be made locally in MFBs in intact slices in 1 hour, whereas somatic Halo-actin requires a longer time to be trafficked into MFBs of the CA3b region.

**Figure 2.**
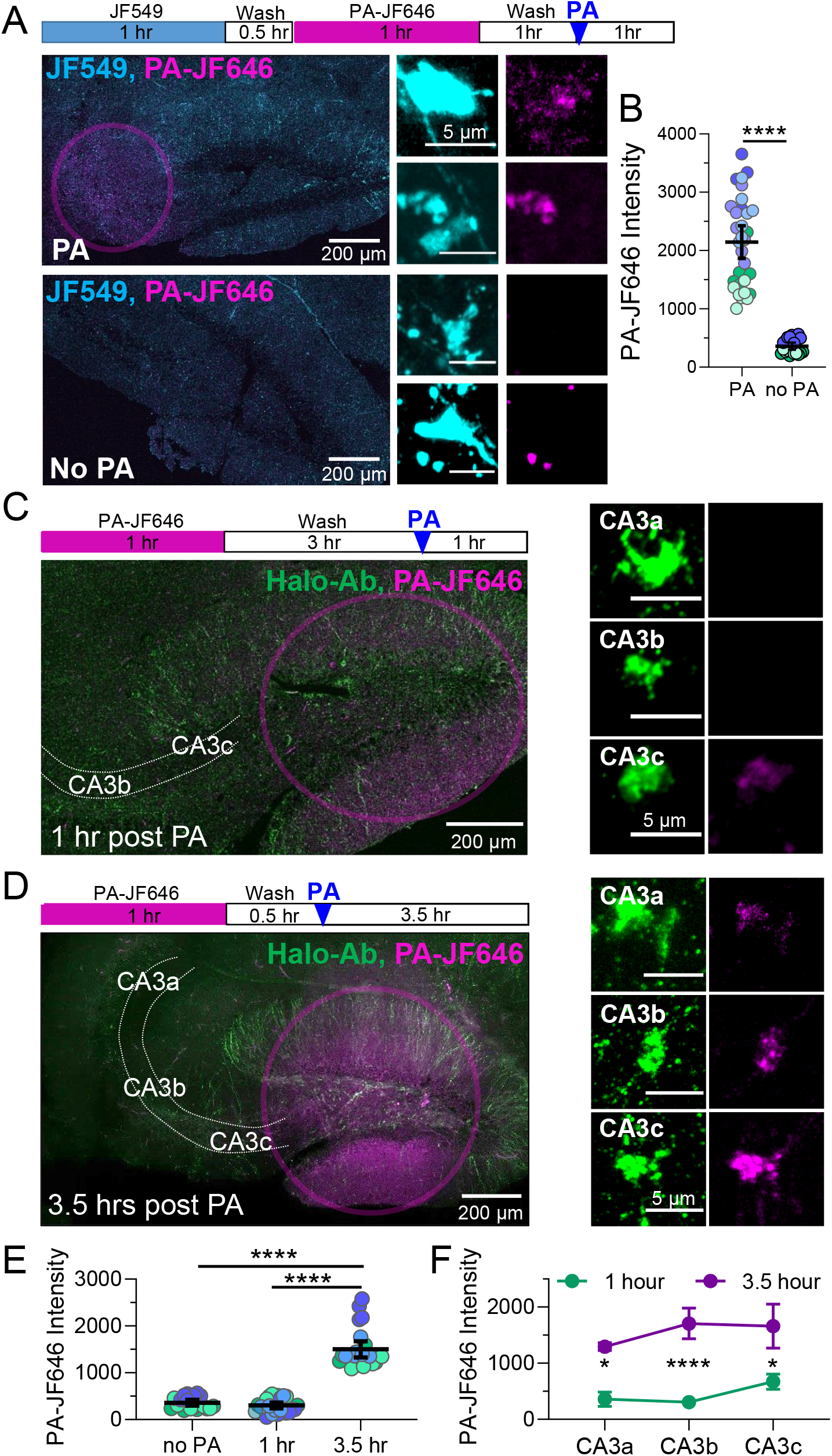
Locally synthesized actin contributes to MFB actin pool within 1 hour. C. (*Top*) Timeline of Halo-actin pulse-chase photoactivation (PA) experiment. (bottom left) Representative high-res Airyscan confocal widefield images of slices from Halo-actin injected mice which were incubated with JF549, followed by PA-JF646, then photoactivated in stratum lucidum (pink circle) or no PA for controls. (center) MFBs from slices that underwent PA had significantly greater levels of newly synthesized Halo-actin than those with no PA (scale bar represents 5 μm). D. Quantification of locally synthesized Halo-actin measured by intensity of PA-JF646 inside MFBs identified using JF549. PA: 2144 ± 137.2 v. no PA: 359.7 ± 29.2 (Mean ± S.E.M.); Mann-Whitney, U = 0, p < 0.0001. n = 31, 20 boutons, 5,3 slices respectively, 3 animals. For all Superplots in this figure, black line and bar represent the mean ± 95 % confidence interval. Points representing individual boutons are color-coded by slice. E. (*Left, top*) Timeline of 1 hour post Halo-actin PA experiment. (*Left, bottom*)Representative Airyscan confocal widefield images of slices from Halo-actin injected mice which were photoactivated in DG (pink circle) at 1 hour prior to fixation. (*Right*)Representative high-res Airyscan confocal images of MFBs with all Halo-actin marked using Halo-Ab staining at different distances from the DG somas (CA3a, CA3b, CA3c) 1 hour after PA (scale bar represents 5 μm). F. (*Left, top*) Timeline of 3.5 hour post Halo-actin PA experiment. (*Left, bottom*)Representative Airyscan confocal widefield images of slices from Halo-actin injected mice which were photoactivated in DG (pink circle) at 3.5 hours prior to fixation. (right) Representative high-res Airyscan confocal images of MFBs with all Halo-actin labeled using Halo-Ab staining at different distances from the DG somas (CA3a, CA3b, CA3c) 3.5 hours after PA (scale bar represents 5 μm). G. Quantification of PA-JF646 mean intensity in single MFBs revealed that there was no significant difference between Halo-actin levels slices with no PA vs slices that were fixed 1 hour after PA in the CA3b region where electrophysiological recordings of MF-LTP are performed. After 3.5 hours, PA-Halo-actin from DG is detectable in CA3b MFBs. No PA: 359.7 ± 29.20 v. 1 hour: 300.5 ± 30.95 v. 3.5 hours: 1501 ± 84.0 (Mean ± S.E.M.); One-Way ANOVA with Tukey’s test for Multiple Comparisons, F [2,65] = 147.3, p < 0.0001. n = 20, 24, 24 boutons respectively, 3-4 animals. H. Summary data showing mean intensity averaged by slice demonstrates significant increases in all CA3 subregions 3.5 hours v. 1 hour after PA. Two-Way ANOVA with Sidak’s Test for Multiple Comparisons, F[1,21] = 48.54, p < 0.0001. n = 5 slices (1 hr), 4 slices (3.5 hrs), 4 animals.

### MFBs contain ribosomes

Given our data showing local synthesis of actin, we next examined if the protein synthesis machinery was present in MFBs. Resolving ribosomes in presynaptic terminals is challenging with conventional antibody labeling, therefore we utilized a cell-specific strategy. The Ribotag mouse has loxP sites flanking the exon 4 of the large subunit ribosomal protein 22 (RPL22) genomic loci enabling Cre recombinase induced expression of a hemagglutinin-tagged RPL22^HA^ (Sanz et al., 2009; Shigeoka et al., 2016). To specifically target GCs, achieve sparse labeling of MFBs and cell-specific expression of RPL22^HA^, we injected lentivirus encoding ChiEFtom2A-Cre or ChiEFtom2A-GFP under the control of the C1QL2 promoter (Barthet et al., 2018) into the hippocampus of the Ribotag mice, resulting in expression of channelrhodopsin (ChiEF) fused to tdTomato. The ChiEFtom2A fusion protein provides a membrane delimited MFB label within which we quantified the HA signal. In control GFP-expressing mice, no HA immunolabeling was detectable in the GC soma or axon as expected (**Figure 3A**). In the Cre-expressing mice, RPL22^HA^ labeling was significantly increased in the MFBs above background levels of HA staining in GFP controls (**Figure 3B,C**), strongly suggesting that MFBs contain ribosomes.

**Figure 3.**
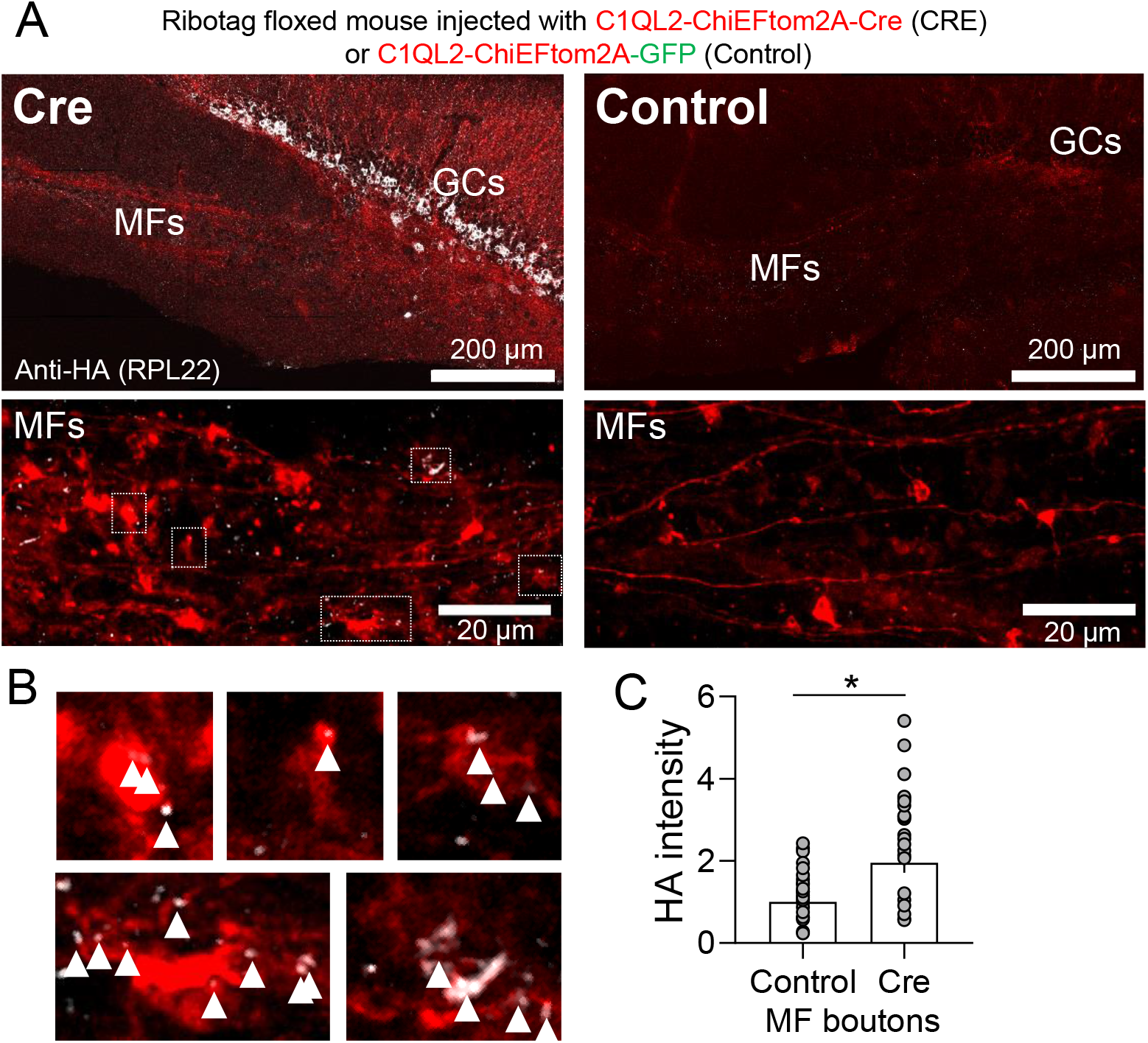
Ribotag mouse reveals MF axons and boutons contain ribosomes. A. (*Top*) Representative widefield image of acute hippocampal slice from Ribotag transgenic mouse injected with lentivirus encoding for ChIEF-tdTomato driven by C1QL2 promoter for dentate GC-specific expression. Note GCs labeled with HA show expected pattern of RPL22 distribution; i.e. excluded from the nucleus. (*Bottom*) MFBs from Cre+ mice show HA labeling in boutons and axons. Boxes indicate boutons magnified in B B. Cre+ MFBs magnified from panel A. White arrowheads indicate RPL22-HA labeling inside contour of bouton. C. Quantification of HA intensity inside MF boutons revealed that Cre expression significantly increased HA intensity above background (Control). Control: 1.0 ± 0.097 v. CRE: 1.95 ± 0.24 (Mean ± S.E.M.); Mann-Whitney, U = 449, p = 0.011. N = 39, 35 boutons, 7, 5 slices respectively, 4 animals.

### MF-LTP involves protein synthesis-dependent structural and functional changes

Given that postsynaptic plasticity is associated with activity-dependent local protein synthesis (Nakahata and Yasuda, 2018), including actin (Yoon *et al*., 2016), we sought to determine whether presynaptic MF-LTP is also associated with local protein synthesis by measuring new Halo-actin synthesis specifically in MFBs as in **Figure 1**. We found that delivering a MF-LTP induction protocol (see Methods) resulted in a modest increase (19% ± 7%) in Halo-actin levels 1 hour later (**Figure 4A,B**), indicating that MF-LTP induces protein synthesis in MFBs. The frequency of boutons with the lowest Halo-actin intensity was reduced in slices that received the LTP induction protocol, suggesting activity may convert MFBs from a translationally quiescent to active state (**Figure 4C**).

**Figure 4.**
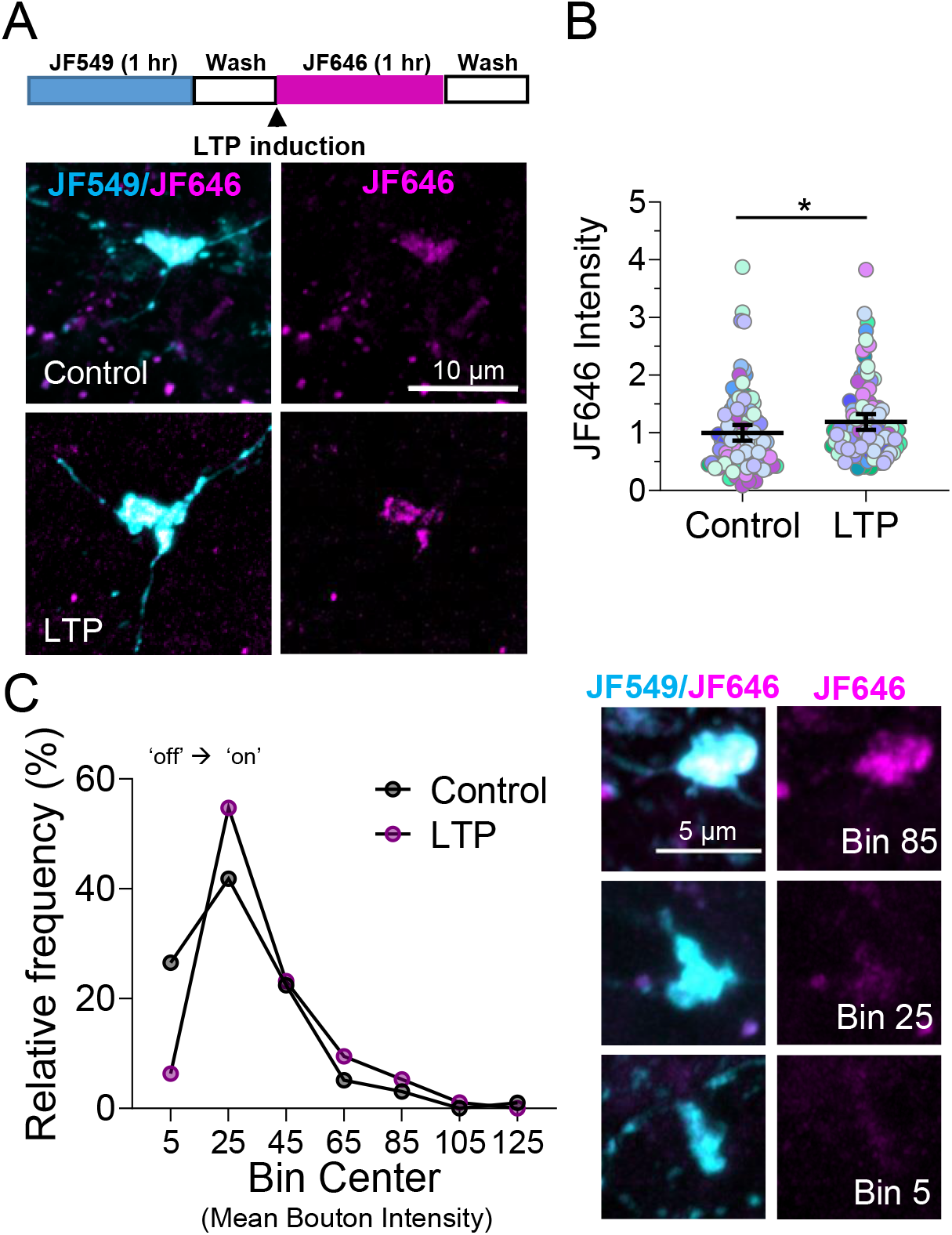
LTP elicits changes in local actin synthesis in MF tract. A. (*Top*) Timeline of Halo-actin LTP experiment. (*Bottom*) Representative high-resolution Airyscan confocal images of pulse-chase labeled mossy fiber boutons in Halo-actin injected slices from Control v. LTP slices. B. MF-LTP increases in newly synthesized Halo-actin in individual MF boutons Control: 1.0 ± 0.07 v. LTP: 1.19 ± 0.07 (Mean ± S.E.M.); Mann-Whitney, U = 3717, p = 0.015, n = 98, 95 boutons respectively, 13,slices, 5 animals. For all Superplots in this figure, black line and bar represent the mean ± 95 % confidence interval. Points representing individual boutons are color-coded by slice and normalized to mean of Control. C. (*Left*) Relative frequency (%) histogram of bouton intensity values indicates LTP primarily impacts low intensity boutons, shifting translationally quiescent boutons (i.e. Bin Center 5) to a translationally active state (i.e. Bin Center ≥ 25). (*Right*) Representative boutons from indicated Bins corresponding to histogram on left.

In order to determine whether MF-LTP-induced changes in Halo-actin levels might be linked to regulation of MFB structure, we investigated whether potential structural changes associated with MF-LTP required protein synthesis. To achieve sparse labeling and optogenetic control over MFBs, we injected the C1QL2-ChiEFtom2A-Cre or -GFP lentivirus as in Figure 3 (**Figure 5A**). Optogenetic activation of GC axons (i.e. MFs) in acute slices (**Figure 5B**) led to increased MFB volume measured 1 hour after LTP induction. Blockade of translation by bath application of 80 μM CHX prevented the MF-LTP-induced increase in bouton volume, but CHX alone had no significant effect on bouton volume (**Figure 5C,D,E**). Notably, CHX application had no effect on basal MF transmission (**Supp. Figure 2A**). MF-LTP and CHX also had no significant effect on filopodial length or number (**Figure 5F,G**).

**Figure 5.**
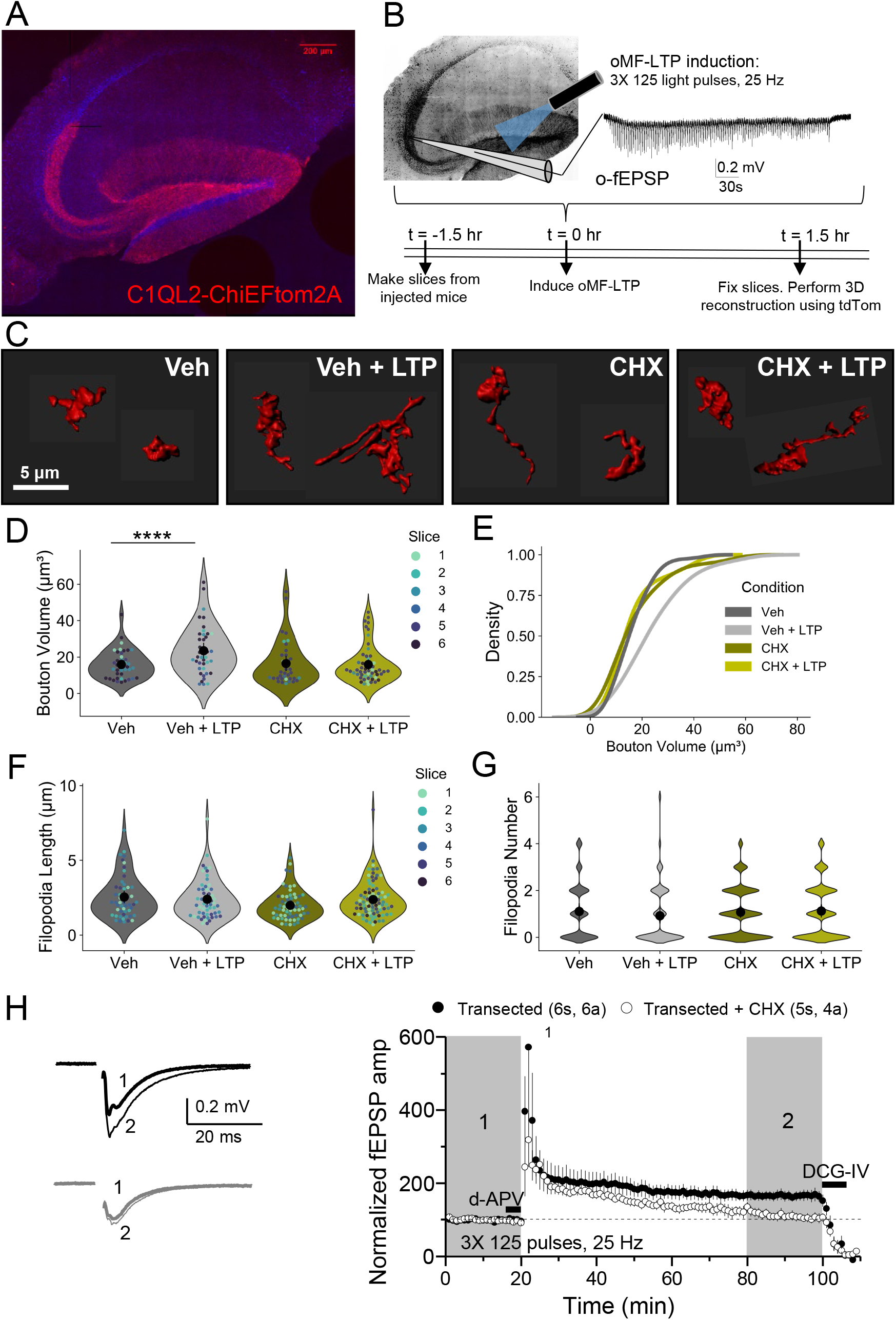
Structural and functional plasticity of MFBs requires protein synthesis. A. Representative widefield image of acute hippocampal slice from mouse injected with lentivirus encoding for ChIEF-tom2A driven by C1QL2 promoter for dentate GC-specific expression. B. Representative optogenetically-induced, extracellular field excitatory postsynaptic potentials (o-fEPSPs) recorded in CA3 resulting from optogenetic light activation (470 nm) in the hilus of slices from A. using MF-LTP induction protocol (125 pulses, 25 Hz, 3x). C. Representative 3D reconstructions of giant MFBs from WT c57 mice injected with C1QL2-ChiEFtom2A-GFP. D. Quantification of MFB volume (μm^3^) from 3D reconstructions reveals MF-LTP induction results in significant increase in bouton volume 1 hour after LTP that is blocked by bath application of protein synthesis inhibitor, cycloheximide (CHX, 80 μM). Veh: 15.94 ± 1.21 v. Veh + LTP: 23.50 ± 2.10 v. CHX: 16.46 ± 2.05 v. CHX + LTP: 15.86 ± 1.36; Two-Way ANOVA with Tukey’s test for Multiple Comparisons F[1,168] = 5.82, p = 0.017, n = 41, 41, 36, 54 boutons respectively, 6 slices, 3 animals. For all Superplots in this figure, individual points representing boutons are color-coded by slice. Large black circle and bar represent the mean ± 95 % confidence interval. E. LTP induction results in significant shift in the distribution of bouton volume when protein synthesis is intact. KS test (*Left*) Z = 3.11, p < 0.00001, (*Right*) Z = 1.2, p = 0.11 F. Length of filopodia is not altered by LTP induction or blockade of protein synthesis. Veh: 1.10 ± 0.18 v. Veh + LTP: 0.91 ± 0.16 v. CHX: 1.10 ± 0.11 v. CHX + LTP: 1.12 ± 0.14; Two-Way ANOVA with Tukey’s test for Multiple Comparisons F[1,252] = 3.00, p = 0.08, n = 52, 57, 64, 83 filopodia respectively, 6 slices, 3 animals. G. Number of filopodia per bouton is not altered by LTP induction or blockade of protein synthesis. Veh: 1.10 ± 0.18 v. Veh + LTP: 0.92 ± 0.16 v. CHX: 1.07 ± 0.12 v. CHX + LTP: 1.12 ± 0.14; Two-Way ANOVA with Tukey’s test for Multiple Comparisons F[1,263] = 0.63, p = 0.43, n = 48, 62, 83, 74 boutons respectively, 6 slices, 3 animals. H. MF-LTP is intact in transected slices, but blocked by bath application of cycloheximide (CHX, 80 μM). Transected: 164.92 ± 13.66 v. transected + CHX: 111.32 ± 11.75; Two sample t-test, p = 0.018, n = 5s, 4a (Transected + CHX) 6s, 6a (Transected).

To test whether functional MF-LTP (i.e. LTP of MF-CA3 synaptic transmission) requires local protein synthesis, we performed recordings in acute WT mouse hippocampal slices where we made a small cut to sever the GC somas from the presynapse (**Figure 5H**), as in Figure 1. Transecting MFs had no effect on the induction or expression of MF-LTP, but MF-LTP was impaired by bath application of the protein synthesis inhibitor CHX (**Figure 5H**). Taken together, these results strongly suggest that MF-LTP requires presynaptic protein synthesis for both synaptic strengthening and enlargement of bouton volume.

### Loss of presynaptic FMRP impairs activity-dependent protein synthesis

MFBs express the mRNA translational repressor FMRP (Christie *et al*., 2009) which has been implicated in local postsynaptic protein synthesis-dependent plasticity (Bagni and Zukin, 2019; Christie *et al*., 2009). We therefore hypothesized that FMRP regulates MF plasticity by modulating local presynaptic protein synthesis. To quantify newly synthesized proteins in MFs, we used a brief pulse of puromycin labeling in acute hippocampal slices to tag nascent peptides (Hafner *et al*., 2019; Schmidt et al., 2009). Inducing MF-LTP in WT mice in the presence of antagonists of ionotropic and metabotropic glutamate receptors (5 μM NBQX, 50 μM d-APV, 4 μM MPEP, 50 μM LY367385) to block synaptic transmission, led to significantly increased puromycin labeling in the MF tract (i.e. stratum lucidum) but not stratum radiatum (**Supp. Figure 3A**), suggesting that the increase in protein synthesis is exclusively presynaptic and not a result of increased postsynaptic activity. As previously reported (David et al., 2012), we confirmed that protein synthesis was blocked by anisomycin, a competitive inhibitor of puromycin that blocks translation elongation by binding to the peptidyl-transferase center (Grollman, 1967), but not CHX, an elongation inhibitor that binds downstream of puromycin in the E site of the ribosome, which resulted in a reduction in puromycin labeling in the MF tract (**Supp. Figure 3B**). In slices from *Fmr1^fl/fl^* mice in which FMRP was conditionally deleted from presynaptic GCs via targeted injection of AAV-CAMKII-mCherry-Cre (cKO), protein synthesis increased under basal conditions and further increased by LTP induction in stratum lucidum where the MFBs reside compared to control slices injected with AAV-CAMKII-mCherry **(Figure 6A,B)**. Importantly, protein synthesis measurements from stratum radiatum, where FMRP expression was intact, were not altered by LTP or cKO of *Fmr1* (**Figure 6C**). For these experiments, AAV was used to achieve higher population of Cre expressing cells to ensure widespread FMRP loss in MFBs (**Supp. Figure 3C**).

**Figure 6.**
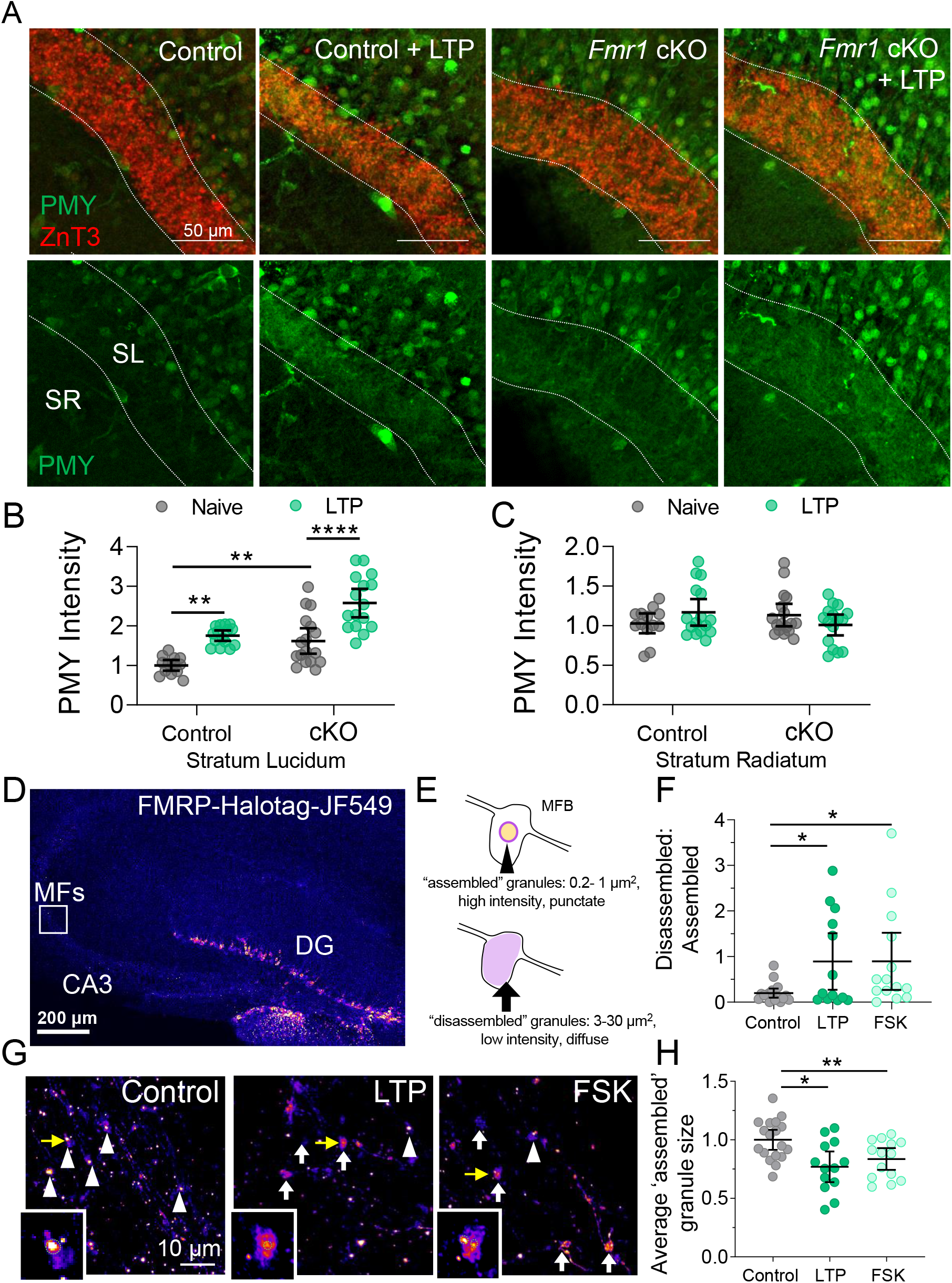
Presynaptic FMRP regulates activity-dependent protein synthesis. A. (*Top*) Representative images of puromycin labeling (PMY) in slices from Fmr1 fl/fl mice injected with CAMKII-mCherry-Cre or CAMKII-mCherry for WT and Fmr1cKO. Dotted lines indicate the MF tract (stratum lucidum, SL) labeled with marker ZnT3. The stratum radiatum (SR) was measured as a control. LTP was induced with electrical stimulation (3X 125 pulses, 25 Hz). B. *Fmr1* cKO caused an increase in PMY labeling in SL and was further increased with LTP induction. WT: 1.0 ± 0.03 v. WT + LTP: 1.75 ± 0.08 v *Fmr1* cKO: 1.62 ± 0.07 v. *Fmr1* cKO + LTP: 2.58 ± 0.06 (Mean ± S.E.M.); Two-Way ANOVA with Tukey’s test for Multiple Comparisons, Row: F[1,57] = 30.94, p < 0.0001, Column: F[1,57] = 43.5, p < 0.0001. N = 13, 15, 17, 16 slices respectively, 8 mice. Points represent individual slices and are normalized to mean of WT. C. *Fmr1* cKO did not increase in PMY labeling in stratum radiatum. WT: 1.0 ± 0.03 v. WT + LTP: 1.17 ± 0.08 v *Fmr1* cKO: 1.13 ± 0.07 v. *Fmr1* cKO + LTP: 1.01 ± 0.06 (Mean ± S.E.M.); Two-Way ANOVA with Tukey’s test for Multiple Comparisons, Interaction: F[1,58] = 3.72, p = 0.06. N = 18,14, 15, 16 slices respectively. Points represent individual slices and are normalized to mean of WT. D. (*Left*) Representative widefield image of acute mouse hippocampal slice expressing FMRP-Halotag fusion protein in GCs. E. Schematic of parameters defining ‘assembled’ granules and ‘disassembled’ granules. F. Ratio of ‘assembled’ to ‘disassembled’ granules is reduced by electrical LTP or chemical LTP induction with forskolin (50 uM, 10 min). Control: 0.197 ± 0.05 v. LTP: 0.891 ± 0.3 v. FSK: 0.894 ± 0.3 (Mean ± S.E.M.); One-Way ANOVA with Tukey’s test for Multiple Comparisons, F[2,42] = 3.828, p = 0.0297; n = 18, 13, 11 slices respectively G. Representative high-res Airyscan confocal images of MFBs expressing Halo-FMRP. High intensity large ‘assembled’ granules in MFBs are indicated with white arrowheads and low intensity, diffuse ‘disassembled’ granules are indicated with white arrows. Insets are granules indicated with yellow arrows. Green dotted line represents contour of ‘assembled’ granule area quantified in panel F. H. LTP induction with electrical stimulation (3X 125 pulses, 25 Hz) or forskolin (50 μM, 10 min) reduced average size of ‘assembled’ granules after 10 minutes. Control: 1.00 ± 0.04 v. LTP: 0.77 ± 0.06 v. FSK: 0.84 ± 0.04 (Mean ± S.E.M.); One-Way ANOVA with Tukey’s test for Multiple Comparisons, F[2,42] = 6.65, p = 0.003; n = 18, 13, 11 slices respectively.

To determine whether FMRP granules are remodeled by MF activity, we expressed mouse FMRP fused to the Halotag (Halo-FMRP) in GCs. Expression of this construct in hippocampal neuron culture revealed strong colocalization of the Halo-FMRP with endogenous FMRP in neuronal somas and neurites (**Supp Figure 4A**). Upon expressing Halo-FMRP in GCs and labeling it in slices through bath application of JF549-HTL (20 nM) (Grimm *et al*., 2015), we detected FMRP localized to MFBs, in agreement with previous reports (**Figure 6D**) (Christie *et al*., 2009). Assembled axonal FMRP granules, which are reported to include FMRP together with ribosomes and mRNA (Akins *et al*., 2017; Chyung *et al*., 2018), can be identified by high intensity fluorescent puncta with sizes in the range of 0.2-1 μm^2^ in MFBs, whereas sites of granule disassembly were indicated by patches of lower intensity, diffuse puncta with an area between 3 – 30 μm^2^ (**Figure 6E**). By counting the ratio of ‘disassembled’ to ‘assembled’ granules, we found that the MF-LTP induction protocol, as well as transient bath application of the adenylyl cyclase activator forskolin (FSK, 50 μM for 10 min), a manipulation that induces chemical LTP of MF-CA3 transmission (Nicoll and Schmitz, 2005), both led to increased ratio of ‘disassembled’ granules in acute hippocampal slices, suggesting translational activation (**Figure 6F,G**). Moreover, the size of ‘assembled’ FMRP granules was also reduced in both forms of LTP induction (**Figure 6G,H**) demonstrating the dynamic nature of FMRP granules. Previously, activity-dependent dephosphorylation of FMRP by PP2A had been implicated in FMRP granule disassembly and translational de-repression (Narayanan *et al*., 2007; Tsang et al., 2019). Consistent with this observation, the addition of PP2A inhibitor, okadaic acid (OKA, 25 nM), abolished the reduction in granule size following MF-LTP and FSK application (**Supp. Figure 4B**).

### Loss of FMRP impairs structural and functional MF-LTP

It has been shown that *Fmr1* KO mice exhibit deficits in MFB density (Ivanco and Greenough, 2002; Mineur et al., 2002) and altered presynaptic plasticity at other CNS synapses (Deng et al., 2011; Koga et al., 2015), suggesting that FMRP may be important for regulating activity-dependent structural and functional MF plasticity. FMRP has not been reported to bind β-actin mRNA directly (Ascano et al., 2012; Eliscovich et al., 2017), (but see (Darnell *et al*., 2011). However, FMRP could participate in MFB structural plasticity via indirect regulation of the translatability of β-actin mRNA (Buxbaum *et al*., 2014) or the direct translational regulation of other cytoskeletal proteins (Feuge et al., 2019). To determine if presynaptic FMRP is involved in the regulation of activity-dependent presynaptic structural modifications, we characterized the structural changes associated with MF-LTP in acute hippocampal slices from *Fmr1* cKO mice (Mientjes et al., 2006). We injected *Fmr1* floxed mice with LV-C1QL2-ChiEFtom2A-Cre or LV-C1QL2-ChiEFtom2A-GFP for controls. Although MF-LTP induced an increase in MF bouton volume in control mice, this increase was not seen in *Fmr1* cKO mice (**Figure 7A-C**). Loss of presynaptic FMRP had no effects on filopodial length (**Figure 7D**), but led to a modest, albeit significant decrease in filopodial number in cKO mice following LTP induction (**Figure 7E**). Therefore, FMRP is a critical regulator of presynaptic activity-dependent structural plasticity.

**Figure 7.**
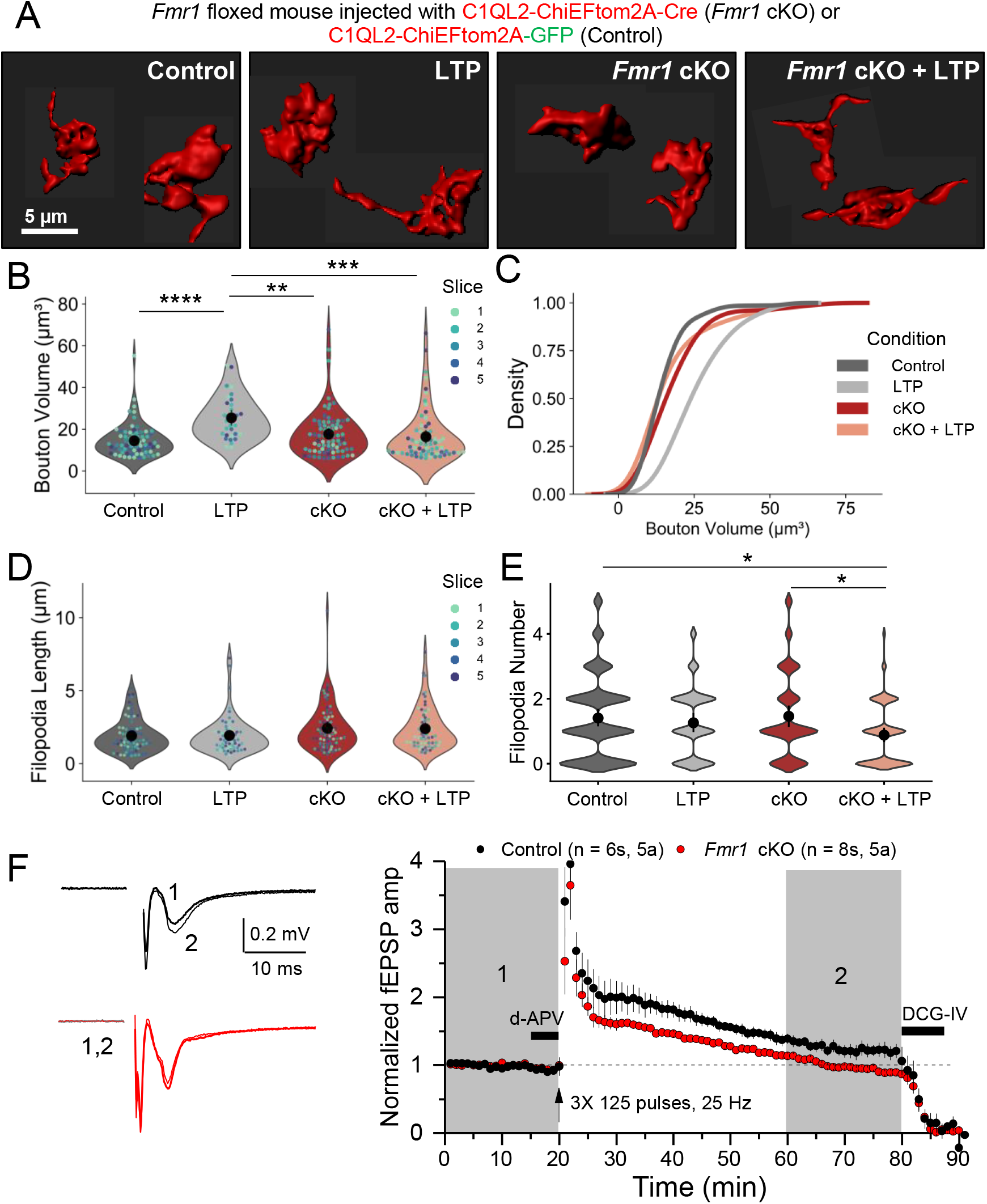
Conditional KO of presynaptic FMRP impairs structural and functional MF plasticity. A. Representative 3D reconstructions of giant MF boutons from *Fmr1*^fl/fl^ mice injected with lentivirus encoding C1QL2-ChiEFtom2A-Cre (*Fmr1* cKO) or C1QL2-ChiEFtom2A-GFP (Control). B. MF bouton volume is increased by LTP induction in WT mice but not in FMRP cKO. Control: 14.87 ± 0.93 v. Control + LTP: 25.44 ± 1.84 v. *Fmr1* cKO: 17.65 ± 1.29 v. *Fmr1* cKO + LTP: 16.38 ± 1.55 (Mean ± S.E.M.). Two-Way ANOVA with Tukey’s test for Multiple Comparisons, F[1,239] = 17.85, p <0.00001; n = 70, 32, 76, 65 boutons respectively, 5 slices, 5 animals. For all Superplots in this figure, individual points representing boutons are color-coded by slice. Large black circle and bar represent the mean ± 95 % confidence interval. C. LTP induction results in significant shift in the distribution of bouton volume. KS test (*Left*) Z = 1.8, p = 0.002, (*Right*) Z = 0.7, p = 0.72. D. Length of filopodia is not altered by LTP induction or absence of presynaptic FMRP. Control: 1.92 ± 0.13 v. Control + LTP: 1.93 ± 0.16 v. *Fmr1* cKO: 2.42 ± 0.18 v. *Fmr1* cKO + LTP: 2.39 ± 0.18 (Mean ± S.E.M.). Two-Way ANOVA with Tukey’s test for Multiple Comparisons, Interaction: F[1.268] = 0.022, p = 0.88, Genotype: F[1.268] = 8.58, p = 0.004; n = 77, 61, 72, 62 filopodia respectively, 5 slices, 5 animals. E. Number of filopodia per bouton is reduced in cKO-LTP. WT: 1.25 ± 0.12 v. WT-LTP: 1.25 ± 0.14 v. cKO: 1.46 ± 0.15 v. cKO-LTP: 0.88 ± 0.11 (Mean ± S.E.M.). Two-Way ANOVA with Tukey’s test for Multiple Comparisons Interaction: F[1,306] = 2.5, p = 0.11, Treatment: F[1,306] = 7.057, p = 0.008; n = 101, 58, 76, 75 boutons respectively, 5 slices, 5 animals. F. (*Left*) Representative EPSPs from extracellular field recording pre- and post-MF-LTP from acute hippocampal slices of *Fmr1*^fl/fl^ mice injected with AAV5-CAMKII-mCherry-Cre or AAV5-CAMKII-mCherry to KO presynaptic FMRP. (*Right*) Summary time-course plot of MF-LTP recordings show MF-LTP is blocked by presynaptic FMRP KO. WT: 124.66 ± 8.47 v. cKO 99.17 ± 1.92 (Mean ± S.E.M.). Two-sample t-test, p = 0.006.

Using extracellular field recordings of MF-CA3 transmission, we found that loss of presynaptic FMRP significantly impaired MF-LTP in cKO slices compared to WT controls (**Figure 7F**). For these experiments, we used an AAV to achieve higher population of Cre expressing cells and thereby greater *Fmr1* KO as we did in Figure 6A. Together, these data not only support a presynaptic localization of FMRP, but also indicate that FMRP granules disassemble during MF-LTP. Moreover, presynaptic FMRP is critical for basal and activity-dependent protein synthesis in the MF tract and can regulate functional MF plasticity.

### Enriched environment alters the functional and structural properties of MFs

Previous work demonstrated that enriched environment (EE), an *in vivo* manipulation known to alter the activity of GCs, alters MF structural plasticity (Caroni et al., 2012; Galimberti *et al*., 2006; Gogolla *et al*., 2009). Given our findings that in vitro manipulation of MF activity altered bouton volume and protein synthesis, we decided to test whether EE can impact the structure and protein synthesis in MFBs in an FMRP-dependent manner. We exposed mice to EE in a large cage with toys and running wheels (**Supp. Figure 5A**) starting at P21 until P36-46 (**Figure 8A**), similar to Gogolla et al., 2009, and compared them to the mice in our Home cage (HC) that were housed in standard cages. Unlike with the induction of MF-LTP, EE did not lead to changes in MF bouton volume compared to HC controls (**Figure 8B-D**), consistent with a previous study (Gogolla *et al*., 2009). However, MFBs from EE mice had significantly longer filopodia with no significant change in the number of filopodia/bouton (**Figure 8E,F**). Thus, EE induces specific structural changes in MFBs that are distinct from those induced by MF-LTP. We also measured gross anatomical features of the MF tract, and found that the infrapyramidal bundle of MFs was significantly longer in the EE exposed animals, as reported previously (Romer et al., 2011), but there was no change in the width or length of stratum lucidum (i.e. the suprapyramidal bundle) (**Supp. Figure 5B**).

**Figure 8.**
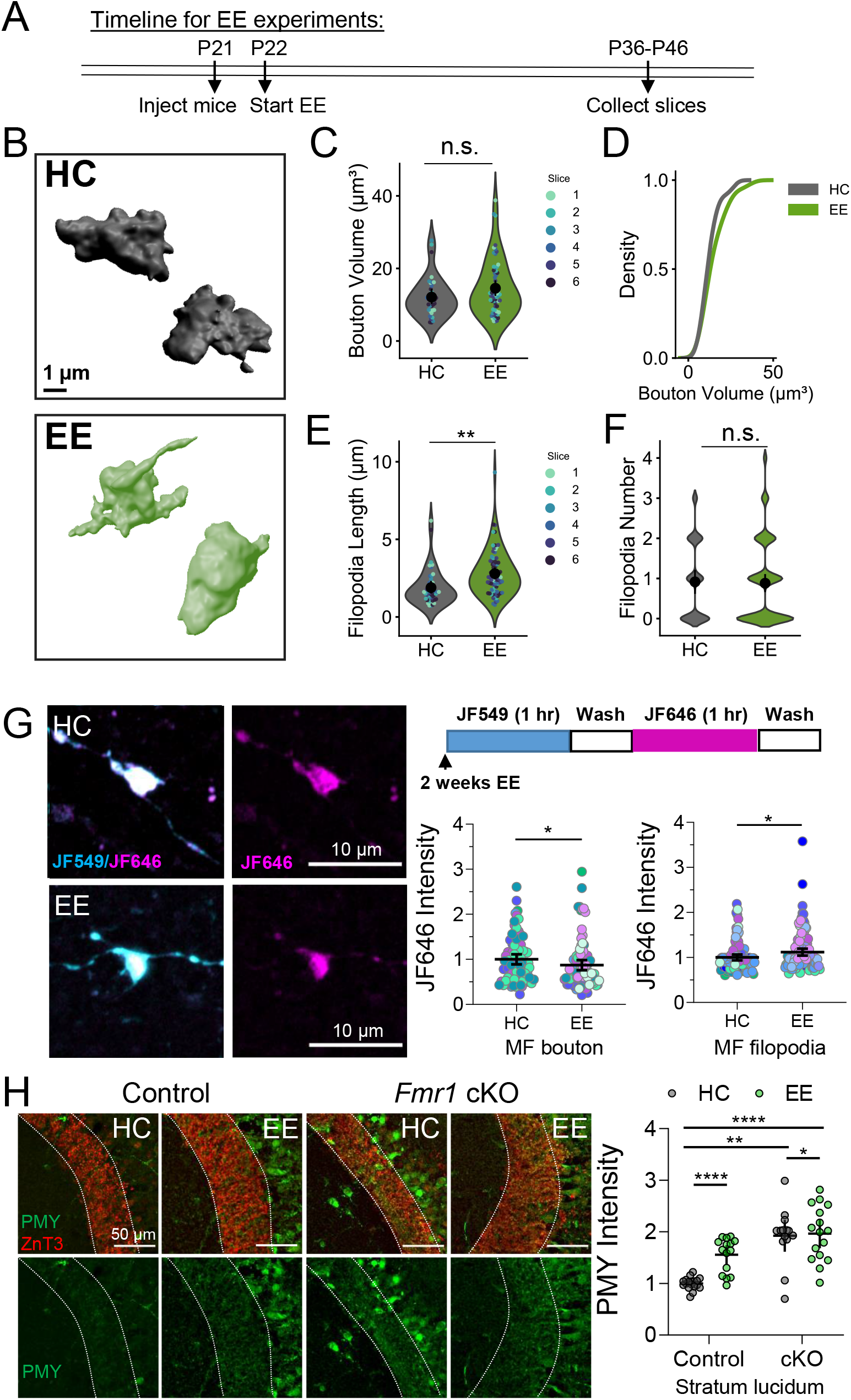
Enriched environment alters structural properties and local protein synthesis of MFs. A. Timeline of enriched environment (EE) exposure. B. Representative 3D reconstructions of giant MF boutons. C. Quantification of MF bouton volume from 3D reconstructions reveals EE causes no change in the overall volume of MF boutons. HC: 13.32 ± 0.75 v. EE: 13.77 ± 0.57(Mean ± S.E.M.); Mann Whitney, Z = −0.72, p = 0.47; n = 97, 158 boutons respectively, 5 animals. For all Superplots in this figure, individual points representing boutons are color-coded by slice. Large black circle and bar represent the mean ± 95 % confidence interval. D. EE does not alter the distribution of MF bouton volume. KS test, Z =0.67, p = 0.79 E. EE significantly changes the length of filopodia. HC: 1.79 ± 0.15 v. EE: 2.70 ± 0.14 (Mean ± S.E.M.); Mann Whitney, Z = −4.41, p = 0.000011; n = 42, 80 boutons respectively, 5 animals. F. EE has no effect on the number of filopodia/ bouton. HC: 0.91 ± 0.14 v. EE: 0.88 ± 0.11; Mann Whitney, Z = 0.34, p =0.73;; n = 47, 92 filopodia respectively, 5 animals. G. (left) Representative images of MF boutons expressing Halo-actin virus from mice in HC or 2 weeks EE. (right, top) Timeline of Halo-actin pulse-chase labeling. (bottom, right) Halo-actin signal measured in MF boutons is reduced in EE mice compared to HC controls. HC: 1.0 ± 0.06 v. EE: 0.87 ± 0.06 (Mean ± S.E.M.); Mann Whitney, Z = 2.19, p = 0.03. (bottom right) Halo-actin signal measured in filopodia of MF boutons is increased in EE mice. HC: 1.0 ± 0.03 v. EE: 1.12 ± 0.04; Mann Whitney, Z = −2.42, p = 0.016. N= 81, 91 boutons/ 100,121 filopodia, 9,11 slices respectively, 5 animals per condition. Points representing individual boutons are normalized to mean of Control. H. Enriched environment did not occlude increased puromycin labeling (PMY) of newly synthesized proteins in MFs of presynaptic cKO of *Fmr1* mice. WT HC: 1.0 ± 0.03 v. *Fmr1* cKO HC: 1.9 ± 0.14 v WT EE: 1.56 ± 0.07 v. *Fmr1* cKO EE: 1.96 ± 0.13 (Mean ± S.E.M.); Two-Way ANOVA with Tukey’s test for Multiple Comparisons, Interaction: F[1,59] = 6.312, p = 0.015. N = 18,14, 15, 16 slices respectively. Points represent individual slices and are normalized to mean of WT HC.

To determine if these experience-induced changes at the MF were associated with changes in protein synthesis, we measured Halo-actin synthesis in individual MFBs (**Figure 8G**). Surprisingly, we found that EE led to a reduction in newly synthesized Halo-actin in boutons but an increase in newly synthesized actin in filopodia of MFBs (**Figure 8G**). These findings are consistent with the lack of change in bouton volume (**Figure 8C**) and the increase in filopodial length (**Figure 8E**). Finally, to assess whether EE resulted in changes in overall newly synthesized protein levels in the MF tract, we performed puromycin labeling as in Figure 6. We discovered that EE led to significant increases in puromycylated nascent peptides in the MF tract (**Figure 8H**). We also tested whether EE altered newly synthesized protein levels in mice with presynaptic cKO of *Fmr1* from GCs. We found that *Fmr1* cKO slices displayed significantly enhanced protein synthesis regardless of EE housing. Together these data indicate that *in vivo* activity of MFs leads to specific changes in presynaptic structure and protein synthesis.

## DISCUSSION

Here, we provide evidence that protein synthesis supports presynaptic structural remodeling associated with presynaptic plasticity. We demonstrate that MFBs can synthesize proteins locally and contain protein synthesis machinery. Moreover, our data demonstrate that protein synthesis in the MF tract is regulated by *in vitro* and *in vivo* activity, which has distinct effects on the structure of MFBs and the local synthesis of a candidate protein, actin. These findings are significant in that protein synthesis is a key mechanism of dendritic spine remodeling associated with postsynaptic forms of plasticity (Nakahata and Yasuda, 2018; Sutton and Schuman, 2006), but its presynaptic function has been heretofore unclear. Along with local protein degradation (Cohen and Ziv, 2017; Monday et al., 2020), local regulation of energy production (Rangaraju et al., 2014), and local structural change, the general acceptance that local protein synthesis occurs in presynaptic compartments supports the notion that presynaptic compartments are discrete, computationally-independent units, akin to dendritic spines.

### Local protein synthesis in excitatory presynaptic boutons

Despite the long-standing dogma that mature mammalian axons and presynaptic terminals do not synthesize proteins, recent evidence to the contrary has amassed, including isolation and identification of presynaptic mRNAs (Akins *et al*., 2017; Hafner *et al*., 2019; Ostroff et al., 2019; Shigeoka *et al*., 2016), visualization of presynaptic ribosomes, translation elongation/initiation factors, and RNA-binding proteins (Akins *et al*., 2017; Hafner *et al*., 2019; Ostroff *et al*., 2019; Scarnati et al., 2018; Shigeoka *et al*., 2016; Younts *et al*., 2016), evidence for altered neurotransmitter release with protein synthesis inhibition (Scarnati *et al*., 2018; Younts *et al*., 2016), and roles for presynaptic protein synthesis in behavior (Ostroff *et al*., 2019) in a wide variety of model systems. Selectively targeting or measuring local protein synthesis in subneuronal compartments is a difficult technical challenge because of the small scale of these structures (Holt *et al*., 2019; Iwasaki and Ingolia, 2017).

We took advantage of the large presynaptic MFBs and used multiple approaches to measure local protein synthesis. First, we optimized the Halo-actin reporter system for broad use in intact brain circuits (Yoon *et al*., 2016). By injecting the Halo-actin reporter virus directly into the DG, we were able to selectively express it in GCs, affording us the spatial resolution to isolate and visualize presynaptic protein synthesis. Using physical transection of MF axons and local photoactivation of JF dyes, we found that local synthesis is a significant contributor to the pool of newly synthesized presynaptic actin. In our slices, the average length of MF axons from the GCs to CA3b subregion was ~ 1 mm. Given published rates of axonal transport of actin are ~2-8 mm/day (Roy, 2020), a protein moving at the maximal velocity would still require ~ 3 hours to reach CA3b from the hilus or GC soma. Consistently, our data (**Figure 2**) suggest that protein trafficking from GC somas/proximal axons to MFBs occurs to some extent after 3.5 hours, but not within 1 hour. Moreover, using cell-specific expression of a tagged ribosomal protein, we validated that the protein synthesis machinery is also present in MFBs.

Even considering multiple mRNAs and polyribosomes, the amount of newly synthesized β-actin likely represents only a small fraction of the large pool of actin at presynapses (Wilhelm et al., 2014), a number that likely varies substantially based on the synapse and preparation. Why then should actin be translated locally? One explanation is that spatiotemporally regulated posttranslational modifications may confer functional distinctions on newly synthesized proteins. It has been proposed that newly synthesized β-actin may be more efficient at nucleation and polymerization of actin filaments (Shestakova et al., 2001), perhaps through fast arginylation at the N-terminus which has been previously shown to increase actin polymerization (Saha et al., 2010), or via chaperone binding to nascent β-actin chains to protect them from glutathionylation, which restricts the rate of polyermization (Shestakova *et al*., 2001). Moreover, the relatively small size of the presynaptic compartment has been suggested to allow the local enrichment of newly synthesized actin monomers, further increasing its ability to create new nucleation sites (Leung *et al*., 2006). Thus, although the contribution of local translation to total subcellular protein content for highly abundant proteins may be small, the functional distinction of somatic versus locally synthesized proteins could be the primary effector of synaptic alterations.

Additionally, we used puromycylation of nascent peptides (Schmidt *et al*., 2009) to study the rate of protein synthesis the MF tract (**Figure 6**). We found that *in vitro* LTP induction enhances overall protein production levels and new actin synthesis in MFs. Moreover, targeted deletion of FMRP increases protein synthesis in the MF tract under basal conditions and more so during high activity (i.e. LTP and EE). We show that EE alone also leads to an enhancement of newly synthesized protein in the MF tract but a reduction in Halo-actin synthesis at the level of the single bouton. To better distinguish activity-induced changes in local protein synthesis in the presynaptic compartment, as opposed to postsynaptic compartments or perisynaptic astrocytic processes (Mazare et al., 2020), excitatory synaptic antagonists were used in order to prevent glutamate-mediated postsynaptic or astrocytic activation. However, we cannot exclude the possibility that other molecules, such as BDNF, may be released from MFs to promote postsynaptic protein synthesis. MF-LTP is induced by a brief activation of GCs specifically (~ 1 minute), whereas EE enhances activity in a wide variety of hippocampal neurons over an extended period of time (2 weeks)(van Praag et al., 2000). EE also increases levels of neurotrophic factors like BDNF and the number of adult-born neurons in the DG (Kempermann et al., 1997). Therefore, homeostatic processes, like changes in inhibitory circuits, formation of new synapses, and/or changes in cellular excitability may potentially occur as a result of EE that could partially overlap with MF-LTP mechanistically.

### Structural plasticity of MFBs

Presynaptic structural plasticity has been reported throughout the brain and can depend on the synapse type, the age of the animal, the size of the bouton, proximity of mitochondria and other organelles (Gogolla et al., 2007; Monday and Castillo, 2017; Smith et al., 2016). At MFs, previous reports suggest structural plasticity is maintained by activity and synaptic remodeling may rely on differential activity among neighboring synapses where larger boutons undergo less structural remodeling (Chierzi *et al*., 2012; Maruo *et al*., 2016). While other studies looked at MF structural changes resulting from diverse forms of robust activity (De Paola *et al*., 2003; Maruo *et al*., 2016; Zhao *et al*., 2012), here we used repetitive burst stimulation of MF axons at 25 Hz (Ben-Simon et al., 2015; Castillo et al., 2002; Kaeser-Woo et al., 2013), to approximate sparse in vivo GC burst firing (10-50 Hz) (Diamantaki et al., 2016; Henze et al., 2002). To our knowledge, our study is also the first to demonstrate that this LTP-induced enhancement in MF bouton volume is dependent on protein synthesis.

At the molecular level, structural changes in MFBs may be critical for functional changes in synapse strength by regulating any number of nodes in the neurotransmitter release process. For example, structural plasticity could enable the insertion and unsilencing of new release sites, alter the coupling distance between calcium channels and primed vesicles, change the size of the active zone and the number of docked vesicles at a synapse, and alter clustering of the release machinery(Monday *et al*., 2018). Actin in the presynaptic terminal is involved in scaffolding of proteins, regulates neurotransmitter release and can be dynamically concentrated to the terminal during bouts of high activity (Morales et al., 2000; Sankaranarayanan et al., 2003). Actin dynamics therefore likely account for the structural remodeling and changes in neurotransmitter release we observed upon MF-LTP induction (Cingolani and Goda, 2008).

### The role of presynaptic FMRP

Previous data have suggested that FMRP granules are dynamically regulated by phosphorylation. Dephosphorylation of FMRP is associated with granule disassembly and translational activation (Narayanan *et al*., 2007) whereas phosphorylation is linked to assembled, translationally-silenced granules (Tsang *et al*., 2019). There is evidence that FMRP loss impairs actin dynamics (Feuge *et al*., 2019; Scharkowski et al., 2018) and, although β-actin mRNA itself is not a bonafide FMRP target (Ascano *et al*., 2012; Eliscovich *et al*., 2017), (but see (Darnell *et al*., 2011), FMRP can directly bind to and regulate the translation of mRNAs encoding key actin cytoskeletal proteins, including cofilin1 (cof1) (Feuge *et al*., 2019), profilin1 (PFN1) (Michaelsen-Preusse et al., 2016), members of the WAVE1 complex and more (Ascano *et al*., 2012). Cytoplasmic FMRP interacting protein 1 (CYFIP1) is a key interacting partner of FMRP which regulates its involvement in both local translation and actin polymerization (Napoli et al., 2008). Interestingly, knockdown of FMRP has been shown to reduce the ‘masking’ of β-actin mRNA from the translation machinery perhaps via interaction with other RNA-binding proteins like zipcode-binding protein (ZBP1) that are known to directly bind β-actin mRNA (Buxbaum *et al*., 2014). Our data are consistent with a model wherein activity-dependent changes in presynaptic FMRP function control the localization and translation of key modulators of the actin cytoskeleton (Michaelsen-Preusse et al., 2018) and/or indirectly regulate the translatability of β-actin mRNA itself. Because our study used a conditional KO strategy to remove FMRP specifically from GCs after P21, we can distinguish the role of FMRP in activity-dependent plasticity at mature synapses from its involvement in early development of MF synapses.

Our data show cKO of FMRP from presynaptic GCs increased protein synthesis in MFBs, but did not affect basal MFB volume. Protein synthesis was further increased by LTP in *Fmr1* cKOs and cKO prevented the expression of structural and functional LTP. This observation suggests that MF-LTP expression may depend on having the appropriate amount of protein synthesized locally, as well as the proper distribution of mRNA species, which is likely also impacted by loss of FMRP.

### Significance of local protein synthesis to in vivo MF function

Due to its sparse firing pattern, the DG is ideally suited for its proposed function of pattern separation by orthogonalizing information coming from the entorhinal cortex (Treves et al., 2008). A single GC makes synapses with only 10-18 postsynaptic CA3 pyramidal cell targets (Claiborne *et al*., 1990), but each one can powerfully regulate CA3 output (Evstratova and Toth, 2014; Henze *et al*., 2000). The MF-CA3 synapse is thought to initiate storage of new information while a direct projection from the entorhinal cortex to CA3 pyramidal cells may serve to initiate retrieval of previously stored representations (Treves *et al*., 2008). Studies of rodents exposed to different habitats, in vivo LTP (Escobar et al., 1997), or to spatial learning tasks show expansion of the infrapyramidal bundle (Routtenberg, 2010), consistent with our EE data (**Supp. Figure 5B**). While most of these studies focused on changes in the gross anatomical features of the MF projection, the microscale structural changes like the LTP-associated MFB enlargement we observed may respresent the fine-tuned establishment and encoding of new representations in the hippocampal circuit. Lastly, given that the subgranular zone is a neurogenic niche, and adult born GCs establish new connections in the mature brain, local presynaptic protein synthesis may be especially critical in the functional integration and pathfinding of immature MF synapses (Toni and Schinder, 2015). Future studies should examine structural plasticity and levels of newly synthesized proteins selectively at adult-born MF-CA3 synapse.

## Acknowledgements

We thank all Castillo Lab and Singer Lab members, and Benjamin Hobson for helpful discussions. We thank Dr. Kostantin Dobrenis, Kevin Fisher, and Vladimir Mudragel of the Einstein Neural Cell Engineering and Imaging Core (supported by The Rose F. Kennedy Intellectual Disabilities Research Center) for their advice and assistance with Airyscan confocal microscopy acquisition and analysis. We are grateful to Dr. Mary Kay Lobo University of Maryland, Baltimore County for donating the Ribotag mice, Drs. Gaël Barthet and Christophe Mulle, Université de Bordeaux, for sharing the lentiviral constructs, and the FRAXA foundation for the *Fmr1*^*f*l/fl^ mouse line. This research was supported by the National Institutes of Health: F31MH114431 to HRM, R01-MH125772, R01-NS113600, R01-NS11543, R01-MH116673, and a pilot grant through NICHD U54 HD090260 to PEC, R01-NS083085 to RHS, R21-MH120496 to YJY, and a shared instrument grant (1S10OD25295) to Konstantin Dobrenis.

## Author Contributions

H.R.M. and P.E.C wrote the manuscript and all authors were involved with editing. H.R.M., S.C.K., Y.J.Y. and P.E.C designed experiments. H.R.M. performed electrophysiology, structural plasticity, and Halo-actin/FMRP experiments. S.C.K performed puromycylation experiments and assisted with structural plasticity and Halo-actin experiments. YJY designed and validated viral constructs.

The authors declare no competing financial interests.

## METHODS

### Animals

All animals were group housed in a standard 12 hr light/12 hr dark cycle. Experimental procedures adhered to NIH and Albert Einstein College of Medicine Institutional Animal Care and Use Committee guidelines. Acute transverse slices were prepared from adolescent male and female Sprague Dawley rats (P18-27) (Charles River) and male and female mice (P21-45): C57 BL/6J (Charles River), Fmr1 fl/fl mice (Dr. David Nelson, FRAXA), RiboTag mice (Dr. Mary Kay Lobo, University of Maryland, Baltimore County).

### Plasmids and Lentiviral production

High-titer lentiviruses were produced in the Einstein Genetic Engineering and Gene Therapy core according to standard protocol. Titer was quantified using fluorescence and ELISA. Mouse Halo-actin construct is described in Halo-actin section below and previously (Yoon *et al*., 2016). Mouse Halo-FMRP construct was generated by replacing the EGFP with the HaloTag coding sequence in the p-EGFP-C1-Flag-mFmr1(wt) vector which was a gift from Stephanie Ceman (Addgene plasmid #87929; http://n2t.net/addgene:87929; RRID:Addgene_87929). The coding sequence of Halo-FMRP was then inserted into the multicloning site of a third generation lentivirus expression vector under the control of the human Synapsin promoter. C1QL2-ChiEFtom2A-GFP or C1QL2-ChiEFtom2A-Cre plasmids were a gift of Drs. Gaël Barthet and Christophe Mulle, Université de Bordeaux, and lentiviruses were made at the Einstein Genetic Engineering and Gene Therapy Core.

### Slice Preparation and Electrophysiological Recording

Acute transverse slices were prepared as follows: briefly, mice were decapitated, and brains were removed quickly and put into ice cold sucrose cutting solution or NMDG cutting solution. The sucrose cutting solution contained (in mM): 215 sucrose, 20 glucose, 26 NaHCO_3_, 4 MgCl_2_, 4 MgSO_4_, 1.6 NaH_2_PO_4_, 2.5 KCl, and 1 CaCl_2_. The NMDG cutting solution contained (in mM): 93 N-Methyl-d-glucamin, 2.5 KCl, 1.25 NaH_2_PO_4_, 30 NaHCO_3_, 20 HEPES, 25 D-glucose, 2 Thiourea, 5 Na-Ascorbate, 3 Na-Pyruvate, 0.5 CaCl_2_, 10 MgCl_2_. Mice over P35 were cut in NMDG. The hippocampi were isolated and cut using a VT1200s microsclicer (Leica Microsystems Co.) at a thickness of 300 μm. These slices were then transferred to 32°C ACSF for 30 min and then kept at room temperature for at least 1h before recording. The artificial cerebral spinal fluid (ACSF) recording solution contained (in mM): 124 NaCl, 26 NaHCO_3_, 10 glucose, 2.5 KCl, 1 NaH_2_PO_4_, 2.5 CaCl_2_, and 1.3 MgSO_4_. Mice under P35 were cut in sucrose solution. After ice-cold cutting, slices recovered at RT (in 50% sucrose, 50% ACSF) for <30 min and then at room temperature (RT) for 1 hr in ACSF. All solutions were bubbled with 95% O_2_ and 5% CO_2_ for at least 30 min. All experiments in acute slices and recordings were performed at 25.5 ± 0.1°C.

For extracellular field recordings, two borosilicate glass stimulating pipettes filled with ACSF were placed in the dentate GC layer at the border of the hilus and a glass recording pipette filled with 1M NaCl was placed in CA3 in *stratum lucidum*. To elicit synaptic responses, paired, monopolar square-wave voltage or current pulses (100–200 μs pulse width) were delivered through a stimulus isolator (Isoflex, AMPI) connected to a broken tip (~10–20 μm) stimulating patch-type micropipette filled with ACSF. Stimulus intensity was adjusted to give comparable magnitude synaptic responses across experiments (~0.2-0.6 mV). Baseline and post-induction LTP synaptic responses were monitored at 0.05 Hz. Stimulation and acquisition were controlled with IgorPro 7 (Wavemetrics). Shaded boxes in figures correspond to when plasticity was analyzed with respect to baseline and when representative traces were collected and averaged. Summary data (i.e. time-course plots and bar graphs) are presented as mean ± standard error of mean (SEM). LTP was triggered by delivering 125 pulses(25 Hz, 3X) in the presence of d-APV (Ben-Simon *et al*., 2015; Castillo *et al*., 2002; Kaeser-Woo *et al*., 2013). At the end of all recordings 1 μM DCG-IV was added to confirm the recording was from a mossy fiber synapse. Only recordings displaying >80% reduction in transmission following DCG-IV application were included.

For recordings in Figure 7, P21-24 *Fmr1*^fl/fl^ mice were injected with an AAV encoding CAMKII-mCherry or CAMKII-mCherry-Cre into the DG using established coordinates (−2.2 posterior to Bregma, 2.0 laterally, and 2.0 ventral from dura) and a total volume of 0.5 μL/ hemisphere at a flow rate of 0.1 μL/min obtained from UNC Vector Core. 2 weeks after injection, acute hippocampal slices were made as described above. After slicing in NMDG, slices were checked for expression using epifluorescence, and excluded if any cells in CA3 were infected. LTP was induced as described above using electrical stimulation. Slices were fixed after recording and saved for post-hoc immunohistochemistry to confirm KO of FMRP (data not shown).

### Immunohistochemistry

Slices were washed twice in PBS then incubated in blocking buffer (4% goat serum in PBS + 2%BSA + 0.1% Tx-100) for 1 hour at RT. Primary antibodies (anti-FMRP, 1:10, FRAXA; anti-HA, rabbit polyclonal, 1:1000 ab9110 Abcam; anti-ZnT3, rabbit polyclonal, 1:500, Synaptic Systems; anti-puromycin 1:1000, EMD MIllipore) were diluted directly into the antibody buffer (blocking buffer without goat serum) and floating slices were incubated overnight at 4C. After 4 washes with PBS, slices were incubated in secondary antibodies (Invitrogen, Carlsbad, CA, USA) diluted in blocking buffer overnight at 4°C. Slices were washed 5X with PBS, then mounted.

### MF Bouton Structural Analysis

*Fmr1*^fl/fl^ mice (Figure 7) and WT C57BL/6J (Figure 5) were stereotactically injected with high titer lentiviral constructs (C1QL2-ChiEFtom2A-GFP or C1QL2-ChiEFtom2A-Cre) into the DG between P21-P24 using established coordinates (−2.2 posterior to Bregma, 2.0 laterally, and 2.0 ventral from dura) using a total volume of 2 μL/ hemisphere at a flow rate of 0.2 μL/min. 2-3 weeks after injection, 300 μm acute hippocampal slices were prepared as described above. LTP was induced using optogenetics in some experiments. Briefly, pulses of blue light were provided with a 473 nm wavelength laser (OEM laser systems Inc., ~ 25 mw/mm power) aimed at the hilus of slices using an optic fiber (200 μm diameter), collimated and delivered through the microscope objective (40X, 0.8 NA) in a submerged chamber with oxygenated ACSF flowing. Cycloheximide (CHX, 80 μM) was added to the bath for conditions indicated in Figure 5. An LTP protocol (125 pulses, 25 Hz, 3X) was delivered in the presence of d-APV using light pulses of 1 ms duration. Slices were incubated at RT for 1 hour after delivering the LTP protocol in oxygenated ACSF before overnight fixation in 4% PFA in PBS. Slices were rinsed and mounted, except for a few controls that were also processed for immunohistochemistry to confirm MF bouton localization. tdTomato-ChiEF fluorescent signal was imaged with 561 nm laser to assess bouton structure. Images were acquired on a Zeiss LSM 880 with Airyscan using a Plan-Apochromat 63x/1.4 Oil DIC M27 and 1.8X zoom. Images were Airyscan processed prior to analysis. Threshold, laser power, and gain were kept constant for each experiment. Pixel width and height was 0.049 μm and voxel depth was 0.187 μm. Z-stacks were taken throughout *stratum lucidum* at similar depths in order to control for PFA permeation. Imaris 9.2 software was used to reconstruct boutons in 3D using the Surface function. MFBs were individually screened and compared with the z-stack after 3D reconstruction to ensure correct identification. Only boutons that were greater than 5 μm^3^ and did not touch the image border were included. For filopodial analysis, z-stacks were maximum projected in FIJI and all protrusions having greater than 0.3 μm displacement from the main bouton and not visibly connected to an axon were considered, then a line was drawn down the filopodia starting at the middle touching the bouton surface to measure length in 2D. For consistency, filopodial analysis was performed by the same person for all conditions, separate from the person that did imaging. All imaging and analysis was performed blind to treatment group.

### Puromycylation

Acute slices were cut as described in the *Slice Preparation and Electrophysiological Recording* section. Puromycin dihydrochloride was diluted into ACSF from a stock of 50mM. Slices were incubated with puromycin (50 μM) for 15 min, then LTP was induced using electrical stimulation in the DG in the presence of d-APV, and slices were incubated for 15 more minutes after LTP induction. For experiments using synaptic blockers: NBQX, 5 μM; d-APV, 50 μM; MPEP, 4 μM; YM298198, 50 μM were added to slices for the first 15 minutes of puromycin incubation and during LTP. After LTP, slices were returned to ACSF + puromycin for 15 more minutes. For CHX and anisomycin controls (80 or 30 μM respectively), slices were pre-incubated in protein synthesis inhibitors for 25 minutes before puro incubation. Slices were then fixed for 1 hour in 4% PFA, blocked for 30 minutes, and stained according to the immunohistochemistry protocol above and imaged on the Zeiss LSM 880 with Airyscan using a LD LCI Plan-Apochromat 25x/0.8 lmm Korr DIC M27and 1.8X zoom. Images were Airyscan processed prior to analysis. Quantification was perfomed using FIJI by drawing regions of interest in the ZnT3 channel to measure Puromycin flurosence intensity in stratum lucidum in the Puromycin channel. For stratum radiatum, regions of interest distal to the stratum lucidum were measured in the Puromycin channel.

### Halo-FMRP

WT C57BL/6J mice were stereotactically injected with high titer Halo-FMRP lentivirus into the DG between P21-P24 using established coordinates (−2.2 posterior to Bregma, 2.0 laterally, and 2.0 ventral from dura) using a total volume of 1.5 μL/ hemisphere at a flow rate of 0.2 μL/min. After 2-3 weeks of recovery, mice were sacrificed and acute hippocampal slices were prepared as described above. For the detection of Halo-FMRP, cell-permeable Halo-ligand conjugated to a tetraalkylrhodamine derivative (JF549-HTL), numbers indicate wavelength of excitation) was bath-applied in a pulse-chase assay. Slices were labelled with JF549-HTL (100 nM) in ACSF in a chamber oxygenated with 95%O_2_/5% CO_2_ for 1 hour to label all Halo-FMRP. For LTP experiments, electrical LTP was triggered using two borosilicate glass stimulating micropipettes with a broken tip ~ 20 μm filled with ACSF placed in the dentate GC layer at the border of the hilus, and MF-LTP protocol was delivered as described above. For chemical induction of LTP, forskolin (50 μM) was bath applied for 10 minutes. For control experiments using okadaic acid, slices were pre-treated with okadaic acid for 30 min prior to and during LTP induction. 15 minutes after the start of electrical or chemical LTP induction slices were fixed with 4% PFA and mounted. Images were acquired on a Zeiss LSM 880 with Airyscan using a Plan-Apochromat 63x/1.4 Oil DIC M27 and 1.8X zoom. Images were Airyscan processed prior to analysis. Threshold, laser power, and gain were kept constant for each experiment. Pixel width and height was 0.049 μm and voxel depth was 0.187 μm. Analysis was performed on high-resolution images of MFBs using FIJI ‘Analyze particles’ with a set threshold to capture high intensity particles of diameter 0.25 μm or larger. Quantification was performed using FIJI to set a high or low intensity threshold and ‘Analyze’ particules of sizes indicated in Results section (Figure 6C) to distinguish between ‘assembled’ and ‘disassembled’ granules.

For controls in Supp. Figure 4A, dissociated hippocampal neurons in culture were infected with Halo-FMRP at DIV9 and fixed at DIV14. IF was performed using monoclonal FMRP antibody 7G1 (deposited by Warren, S.T. to the Developmental Studies Hybridoma Bank at the University of Iowa, Iowa City, IA, USA) at 1:10 dilution and polyclonal HaloTag antibody (Promega Corp. Madison, WI, USA) at 1:500 dilution and FITC-conjugated donkey anti-mouse secondary antibody and Cy3-conjugated goat anti-rabbit secondary antibody (Invitrogen).

### Halo-actin

WT C57BL/6J mice were stereotactically injected with high titer Halo-actin lentivirus into the DG between P21-P24 using established coordinates (−2.2 posterior to Bregma, 2.0 laterally, and 2.0 ventral from dura) using a total volume of 1.5 μL/ hemisphere at a flow rate of 0.2 μL/min. The Halo-actin reporter construct includes FLAG-tag and HaloTag sequences upstream and in-frame of the β-actin coding sequence followed by the 3’ UTR of β-actin and the nonrepetitive MBSV5 aptamers. After 2-3 weeks of recovery, mice were sacrificed and acute hippocampal slices were prepared as described above. For the detection of newly translated proteins, cell-permeable Halo-ligand conjugated to a tetraalkylrhodamine derivatives (JF549-HTL and JF646HTL; Janelia Research Campus, Ashburn, VA, USA) were bath-applied in the pulse-chase assay. Slices were labelled with JF549-HTL (100 nM) in ACSF in a chamber oxygenated with 95%O_2_/5% CO_2_ for 1 hour to label all the preexisting Halo-actin, then washed out in ACSF for 30 minutes. Next, JF646-HTL (200 nM) was bath-applied to label the newly synthesized Halo-actin, then washed out for 30 min in ACSF. For some controls, the order of the dyes was reversed. The translation inhibitor CHX (80 μM; Tocris) were added to the pulse and included in all subsequent steps until the last washout step before imaging. LTP was induced in a subset of slices using electrical stimulation with 2 glass pipettes in the DG, as described in the section on electrophysiology, in the presence of d-APV.

For experiments involving photo-activation, bath-application of PA-JF646 was performed as described above at 2 time intervals (Figure 2). White light from the Sola Lumencor light engine (Lumencor) was directed throught a 40X objective to focus the photoactivation on a small portion of the tissue. To activate, a mirror was used to reflect the full light spectrum of the laser at the tissue for 1 s. The slices were fixed with 4% PFA, mounted and imaged on a Zeiss LSM 880 with Airyscan using a Plan-Apochromat 63x/1.4 Oil DIC M27 and 1.8X zoom. Images were Airyscan processed prior to analysis. Threshold, laser power, and gain were kept constant for each experiment. Pixel width and height was 0.049 μm and voxel depth was 0.187 μm. Z-stacks were taken throughout *stratum lucidum* at similar depths such that boutons were captured in their entirety (i.e. not cut off). Widefield images were obtained with a Plan-Apochromat 10x/0.45 M27 objective with 1.8X zoom. Analysis was performed using FIJI by hand-drawing an ROI around MFBs based on the pre-existing actin channel, JF-549 or Halo-tag antibody (1:500, Promega, mouse monoclonal) and measuring the newly synthesized actin (in most cases, JF646) within this ROI.

### RiboTag

RiboTag mice were injected with C1QL2-ChiEFtom2A-GFP or C1QL2-ChiEFtom2A-Cre at age P21 and sacrificed after 3 weeks. Animals were transcardially perfused with 4% PFA in 0.1M sodium phosphate buffer (PBS). After 48h fixation in 4% PFA, 125 μm-thick brain coronal sections were prepared using a DSK Microslicer (DTK-1000). Brain slices were kept in PBS until staining as described above in Immunohistochemistry using Rb pAb to HA tag (1:1000, Abcam ab9110). For image quantification in FIJI after Airyscan processing, HA signal that overlapped with tdTomato labeled MFBs was measured in individual boutons.

### Enriched Environment

Home cages (HCs) are standard mouse cages with single housed mice. Their dimensions are 28 cm (11”) x 18 cm (7”), and they have a wire feeder lid and a water bottle. Mice assigned to enriched environment (EE) housing are kept in groups of at least 5 in a large 121 cm (4 ft) x 61 cm (2 ft) enriched cage, containing a feeder, water dispenser, several running wheels, as well as plastic tubes, domes and other structures (Supp. Figure 5A). Objects are rearranged every other day to maintain novelty. Female and male mice are never mixed in the same cage, and all males are housed with littermates. EE refers to both the effects of enrichment strictly defined (exploration, novel objects, increased area), as well as the exercise associated with exploration of the environment, and the use of running wheels. For all EE experiments, one HC and one EE mouse was cut per day, and experimenters were blind to condition for imaging and analysis. Experiments to quantify structure, puromycin labeling, and Halo-actin expression were all performed as described above.

### Data analysis, Statistics and Graphing

Analysis and statistics were carried out in OriginPro 2015 (OriginLab) and Graphpad Prism 7.02. Significance (p <0.05) was assessed with one-way ANOVA (means comparison with *post hoc* Bonferroni test), Student’s paired and unpaired t-tests, Wilcoxon matched-pairs signed rank test, Mann Whitney test, or Pearson’s correlation coefficient, as indicated. Python 3.0 was used for Superplots in structural plasticity figures. All experiments were performed in an interleaved fashion and include at least 3 mice. Experimenters were blind to genotype during recording and analysis.

### Reagents

Stock reagents were prepared according to the manufacturer’s recommendation in water, DMSO (<0.01% final volume during experiments), or phosphate buffered saline (PBS), stored at −20°C, and diluted into ACSF or intracellular recording solutions as needed. d-APV was acquired from the NIMH Chemical Synthesis and Drug Supply Program; salts for making cutting solutions, ACSF, puromycin dihydrochloride and DCG-IV from Sigma Aldrich (St. Louis, MO, USA); CHX, MPEP and YM298198 from Tocris Bioscience (Bristol, UK), anisomycin from ALFA Aesar (Ward Hill, MA, USA), okadaic acid and NBQX from Cayman Chemical (Ann Arbor, MI, USA). Reagents were either acutely bath applied or preincubated with slices/cultures, as indicated in Results.

## SUPPLEMENTAL FIGURE LEGENDS

**Supp. Figure 1. (related to Figure 1) MFBs synthesize Halo-actin locally**

A. Representative DIC widefield image of transected acute hippocampal slice showing recording configuration.

B. Ratio of newly synthesized actin (JF646) to pre-existing actin (JF549) is not altered by transection. Control: 0.062 ± 0.007 v. Transected 0.074 ± 0.006 (Mean ± S.E.M.); Mann-Whitney, U = 1182, p = 0.08. n = 23 boutons, 8,7 slices respectively, 4 animals. Black line and bar represent the mean ± 95 % confidence interval. Points representing individual boutons are color-coded by slice and normalized to mean of Control.

C. Reversing the order of the Halo dye does not affect labeling.

**Supp Figure 2. (related to Figure 5) MF-LTP involves protein synthesis**

Electrophysiological recording from acute hippocampal slice demonstrating that CHX has no effect on baseline MF transmission with 1 hour of bath application. CHX: 0.935 ± 0.03; One sample t-test, p = 0.16.

**Supp. Figure 3. (related to Figure 6) Puromycin labeling in WT mice is increased with LTP**

A. LTP induction in the presence of blockers of ionotropic and metabotropic glutamate receptor blockers increases puromycin labeling (PMY) of newly synthesized proteins in MF tract (stratum lucidum) labeled with marker ZnT3, but not stratrum radiatum. (top) Control: 1.0 ± 0.09 v. LTP: 1.45 ± 0.16, Two-sample t-test, p = 0.027, (bottom) Control: 1.0 ± 0.11 v. LTP: 1.0 ± 0.06, Two-sample t-test, p = 0.98. n = 6 slices, 5 animals (Control); 5 slices, 5 animals (LTP).

B. Puromycin labeling is sensitive to inhibitors of protein synthesis, anisomycin and cycloheximide. labeling (PMY) of newly synthesized proteins in MF tract labeled with marker ZnT3. Control: 1.0 ± 0.03 v. ANISO: 0.45 ± 0.10 v. CHX: 1.32 ± 0.05, One Way ANOVA, F[2,11]= 41.18, p < 0.0001, n = 4 slices, 4 animals.

C. Representative widefield images of mCherry viral expression from slices used in puromycin experiments from Control and *Fmr1* cKO mice.

**Supp. Figure 4. (related to Figure 6) Characterization of Halo-FMRP construct and mechanism of FMRP granule regulation by MF-LTP**

A. Endogenous FMRP (*left*) and FMRP-Halotag (*right*) are colocalized in dendrites of cultured hippocampal neurons. Scale bar denotes 5 μm.

B. Activity-dependent FMRP Halotag granule dissembly is regulated by PP2A. Application of okadaic acid (OKA, 25 nM, 30 min) blocked LTP induced decrease in granule size and had no effect on basal granule size. Control: 1.00 ± 0.04 v. LTP + OKA: 0.89 ± 0.09 v. OKA: 1.04 ± 0.10 (Mean ± S.E.M.); One-Way ANOVA with Tukey’s test for Multiple Comparisons, F[2,43] = 0.98, p = 0.38; n = 18, 13, 15 slices respectively.

**Supp Figure 5. (related to Figure 8) EE alters the mossy fiber bundle.**

A. Enriched environment setup. Mice were housed in a large plexiglass enclosure in groups of 5 or more and have access to running wheels, tubes, toys and ample nesting supplies. Objects are rearranged every other day.

B. Histology of mossy fiber tract revealed expansion of the infrapyramidal bundle in EE, while other measures of gross anatomy of MF tract were unaffected. Arrowheads indicate approximate end of IP, brackets show the width of SL and IP. IP Length of IP: HC: 510.2 ± 43.6 v. EE: 672.70 ± 44.5, Two-sample t-test, p = 0.0251. Width of SL: HC: 50.97 ± 2.94 v. EE: 46.0 ± 3.06, Two-sample t-test, p = 0.27. Width of IP: HC: 28.02 ± 2.21 v. EE: 36.37 ± 4.76, Two-sample t-test, p = 0.11.

