## Supplementary figures and images for "Presynaptic FMRP and local protein synthesis support structural and functional plasticity of glutamatergic axon terminals"

### Supplemental Figures

*Supp. Figure 1*

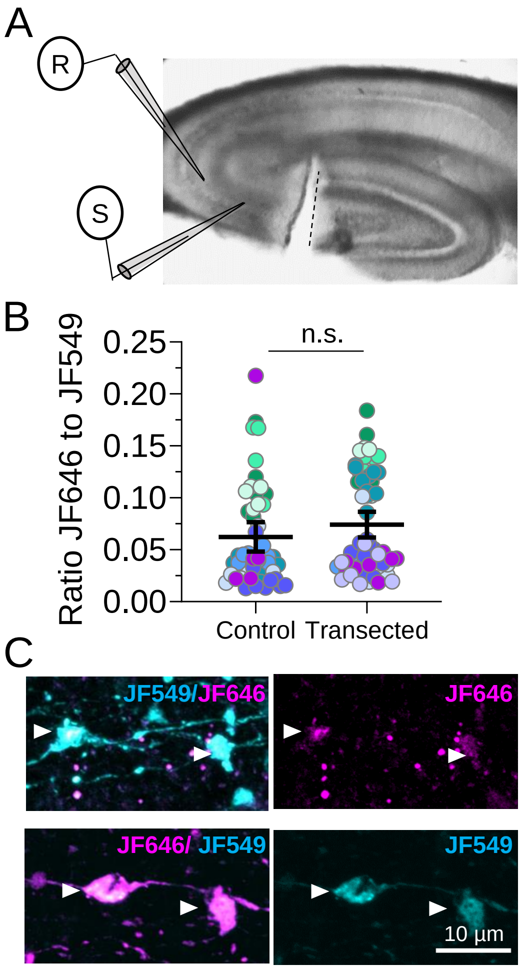

*Supp. Figure 2*

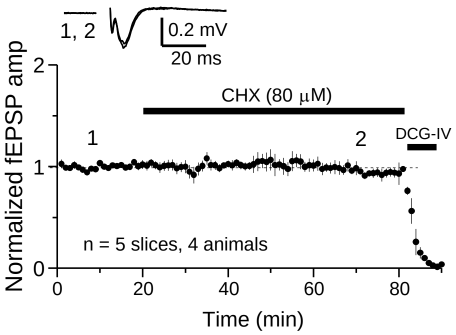

*Supp. Figure 3*

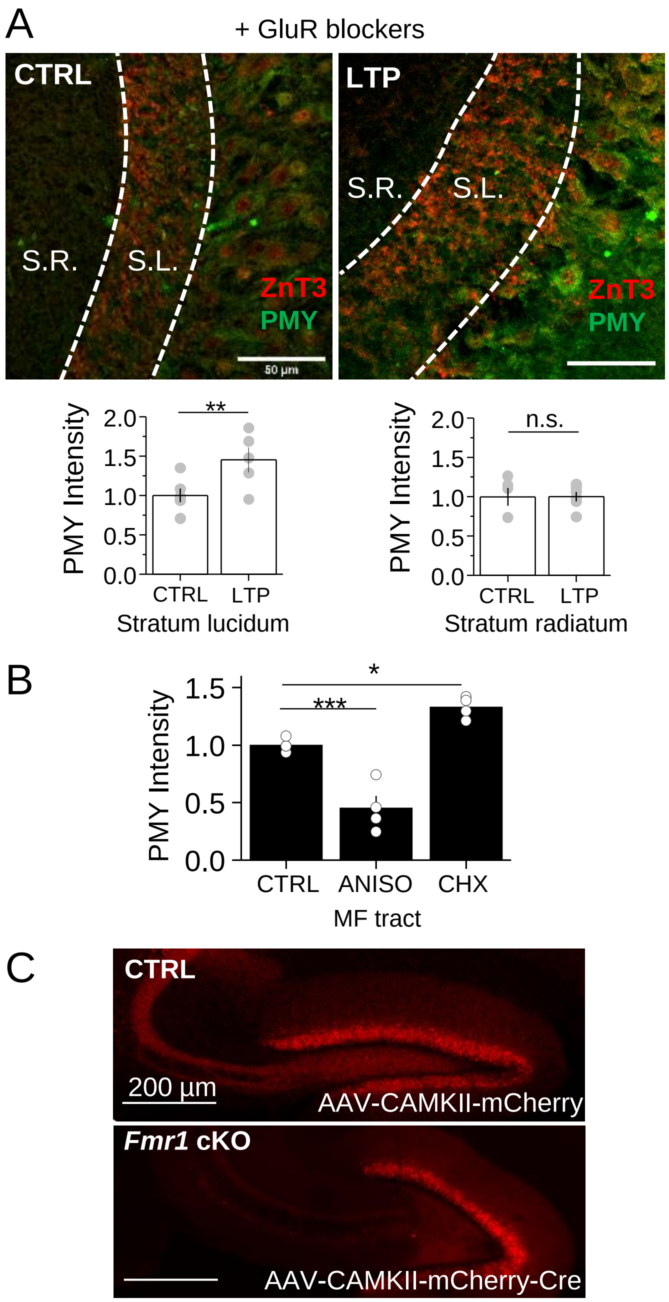

*Supp. Figure 4*

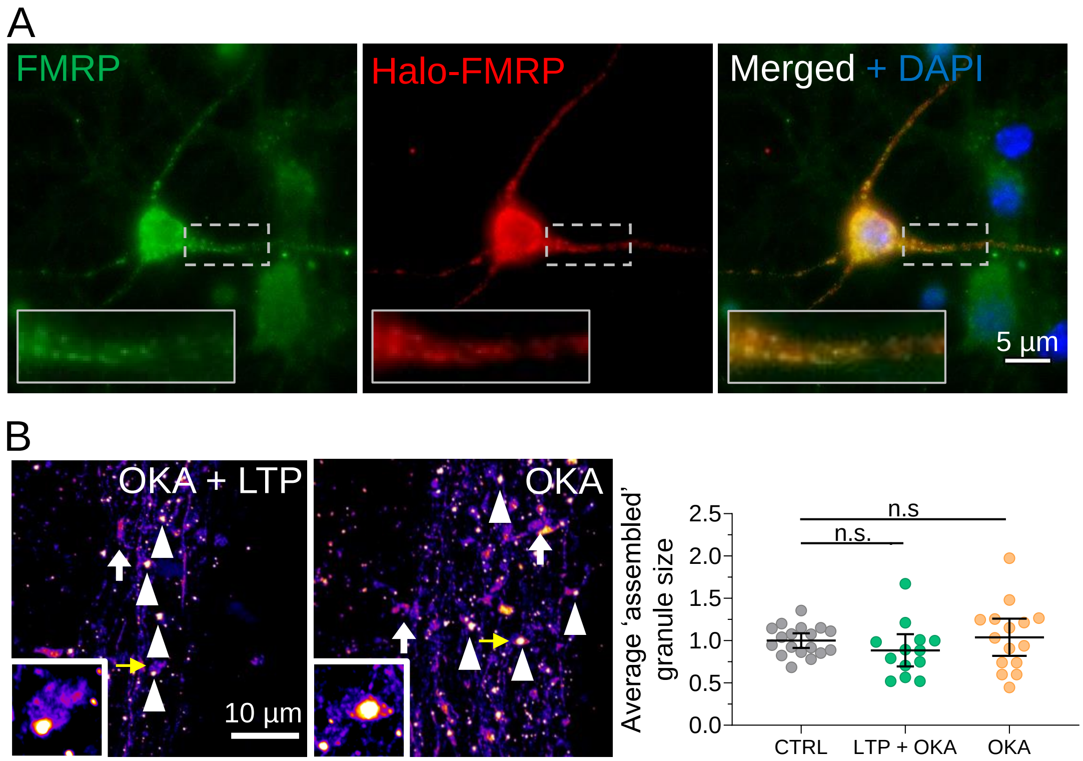

*Supp. Figure 5*

A

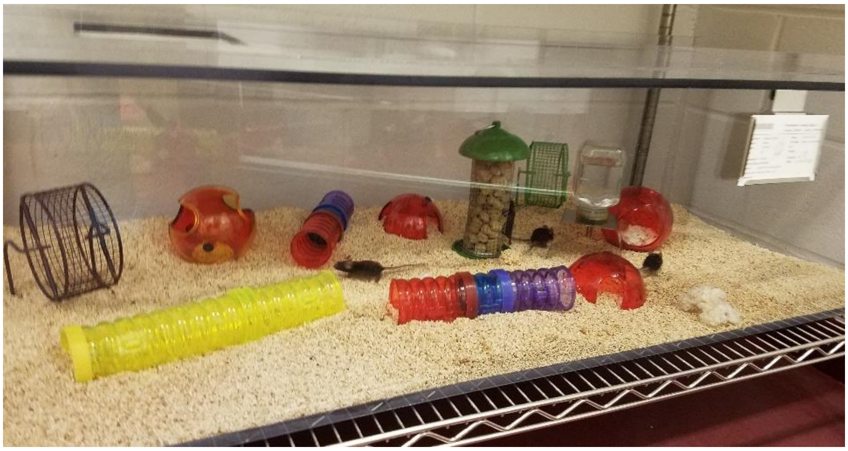

B

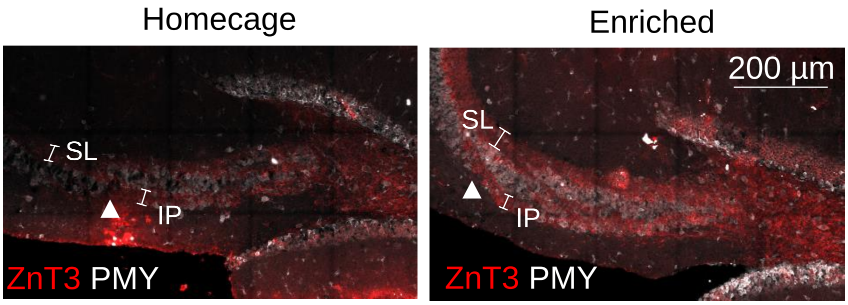


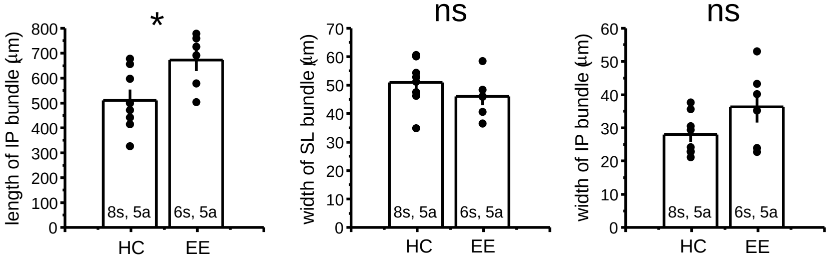
